# Nuclear metabolism oscillation during the cell cycle reveals a link between the phosphatidylinositol pathway and histone methylation

**DOI:** 10.1101/2024.12.20.629614

**Authors:** Antoni Gañez-Zapater, Savvas Kourtis, Lorena Espinar, Laura García-López, Laura Wiegand, Maria Guirola, Frédéric Fontaine, André C Müller, Sara Sdelci

**Affiliations:** Centre for Genomic Regulation (CRG), The Barcelona Institute of Science and Technology, Dr. Aiguader 88, Barcelona 08003, Spain; Universitat Pompeu Fabra (UPF), Barcelona, Spain; CeMM Research Center for Molecular Medicine of the Austrian Academy of Sciences; Vienna, Austria; (now) Charité – Universitätsmedizin Berlin, Corporate Member of Freie Universität Berlin and Humboldt-Universität zu Berlin, and Berlin Institute of Health, Department of Hematology, Oncology, and Cancer Immunology, Berlin, Germany

## Abstract

The progression of the cell cycle is regulated by the expression of specific genes and fluctuations in cellular metabolic states. Previous research has employed cell cycle-based transcriptomics, proteomics, and metabolomics analyses to identify cell cycle-dependent changes at the gene expression, protein, and metabolic levels. However, the role of protein compartmentalization in regulating protein function, coupled with evidence that metabolic enzymes can localize to the nucleus and influence chromatin states, suggests that fluctuations in nuclear metabolism may play a role in regulating cell cycle progression. In this study, we developed an approach to resolve chromatin and nuclear changes during the cell cycle in an unbiased and systematic manner. This was achieved by integrating cell cycle fluorescent reporters with chromatin mass spectrometry and cellular imaging. Our investigation focused on metabolic enzymes and revealed that phosphatidylinositol metabolism localizes to the nucleus in a cell cycle-dependent manner. Moreover, disruption of phosphatidylinositol metabolism affects the nuclear distribution of phosphatidylinositol 4,5-bisphosphate, alters the number and morphology of nucleoli, and influences the maintenance of distinct heterochromatin states throughout the cell cycle. Finally, given the established link between phosphatidylinositol metabolism and methionine synthesis, as well as the differential impact observed on distinct histone marks when phosphatidylinositol metabolism is perturbed, we proposed that distinct pools of methionine may be involved in the maintenance of histone marks that decorate heterochromatin in a cell cycle-dependent manner.

## Introduction

The eukaryotic cell cycle is a highly regulated process that controls cell division and proliferation. Central to this regulatory network is the cell cycle-coordinated control of chromatin, which determines major epigenetic modifications^1^ that fine-tune transcription^2^, control DNA replication^3^ and allow chromosome compaction and correct segregation^4^. Recent advances in live-cell imaging technologies have provided unprecedented insights into the dynamics of cell cycle progression, revealing intricate spatial and temporal patterns of molecular activities^5^. One of such technologies is the Fluorescence Ubiquitination Cell Cycle Indicator (FUCCI) system that enables the visualization of cell cycle phases based on the expression levels of fluorescent gene reporters targeted to specific cell cycle regulators^6,7^. The first generation of FUCCI only allowed the discrimination of proliferating and non-proliferating cells^8,9^. By integrating the tracking of three different cell cycle markers [Chromatin Licensing And DNA Replication Factor 1 (Cdt1), stem-loop binding protein (SLBP) and Geminin] and Histone 1, the more recent FUCCI-4 system allows for real-time monitoring of cell cycle transitions and chromatin condensation during mitosis, revealing insights into the heterogeneity and plasticity of cell cycle progression within populations of cells^10,6^.

Beyond its evident role in chromatin dynamics, emerging evidence suggests that the cell cycle is intricately linked to cellular metabolism^11,12,13^. Metabolic processes, including glycolysis, oxidative phosphorylation and lipid metabolism, provide the necessary energy and building blocks to support DNA duplication, cell growth and proliferation^14,15,16,17,18,19,20,21^. Interestingly, metabolic enzymes have been shown to exert regulatory functions beyond their canonical metabolic roles, influencing signaling pathways and epigenetic modifications that govern chromatin structure and function. Indeed, the nuclear localization of metabolic enzymes regulates chromatin states and fates, influencing histone posttranslational modification^22,23,24,25^ as well as facilitating the DNA damage repair^24,26,27^, regulating transcription^28,29^, and controlling the faithful progression of cell cycle gene expression control^30,31^ and mitosis.

Despite the apparent complexity of the relationship between metabolism and the cell cycle, and the evidence that metabolic enzymes can localize to the nucleus to regulate chromatin states and functions, there has been no attempt to elucidate how nuclear metabolism changes over the cell cycle, nor how these changes affect epigenetic remodeling. In this study, we employed a custom FUCCI reporter to couple cell cycle-based sorting and the isolation of chromatin-bound proteomes, with the objective of identifying metabolic enzymes that fluctuate in their presence on chromatin during the cell cycle. Our results demonstrate that phosphatidylinositol-related enzymes oscillate on chromatin in a cell cycle-dependent manner. This regulation influences the behavior of nucleoli, histone methylation, and the progression of the cell cycle.

## Results

### Optimization and characterization of a FUCCI-3 stable reporter

To characterize the chromatin binding proteomes at the different stages of the cell cycle, a stable U2OS reporter cell line was generated with a customized version of the FUCCI-4 system^32^ henceforth referred to as FUCCI-3. The FUCCI-3 system enabled the tracking of cells through the cell cycle phases using either high-throughput immunofluorescence or fluorescence-activated cell sorting (FACS), thereby facilitating the isolation of cell cycle-specific populations and the quantification of selected markers in a cell cycle-dependent manner (Figure 1A). In addition, since there was no need for cell cycle inhibitors to isolate or track different cell cycle populations, our system avoided the naturally introduced biases due to cell cycle arrest. As with the original FUCCI-4, the FUCCI-3 reporter system expresses fragments of Cdt1, SLBP, and Geminin, respectively tagged with mKO2, Turquoise2, and Clover. However, we eliminated the expression of Histone 1, tagged with the Maroon1 fluorescence marker, which proved to be deleterious to cell proliferation when stably expressed in U2OS cells following lentiviral transduction (Figure S1A). The integration of the expression levels of the Cdt1, SLBP, and Geminin fluorescent markers allowed the classification of cell cycle phases and phase transitions, as previously described^32^. Leveraging this, we developed a pipeline for the analysis of high-throughput microscopy images in the context of the cell cycle (Figure S1B). In brief, images of the FUCCI-3 fluorescent markers were acquired and merged, resulting in the identification of all nuclei. A perinuclear ring region around the nucleus was used to measure the background intensity of the FUCCI fluorescent markers. The background was then subtracted from the mean intensity of the nuclei, allowing for the determination of the corrected level of each of the FUCCI fluorescent markers inside the nucleus. This pipeline enabled a novel microscopy-based analysis of cell cycle, thereby providing a valuable tool for the precise identification and characterization of cell cycle phases at the single nuclei resolution. Removing the Histone 1-Maroon1 reporter allowed the imaging of proteins of interest. In addition, the use of high-throughput microscopy instead of FACS, which is typically used for cell cycle studies, allowed information about cellular structures and subcellular localization of proteins to be preserved. To validate our U2OS FUCCI-3 reporter cell line, we sought to identify cell cycle phases by integrating FUCCI-3 fluorescent markers with nuclear quantification of three well-known cell cycle-associated proteins throughout cell cycle progression: Kiel antigen 67 (Ki67), which is highly expressed in proliferating cells^33^, Cyclin-dependent kinase inhibitor 1A (p21), whose expression promotes cell cycle arrest^34^, and Histone h3 (H3), which is an integral part of nucleosomes and therefore reflects the amount of DNA present in a cell. U2OS FUCCI-3 reporter cells were fixed and stained with antibodies that recognize these markers and analyzed using high-throughput microscopy. The three stained populations (Ki67, p21, and H3) exhibited comparable cell cycle profiles, as illustrated by the FUCCI-3 plots (Figure S1C). In these plots, the intensity of Geminin-Clover is represented on the y-axis, the intensity of the SLBP1-Turquoise2 is represented on the x-axis, and the intensity of Cdt1-mKO2 is shown as a blue-to-red color gradient. The integration of the FUCCI-3 reporters expression enabled the identification of five distinct cell populations. Population 1 was found to correspond to G1 cells, which typically express low levels of Geminin-Clover, low levels of Cdt1-mKO2, and medium levels of SLBP-Turquoise2. Population 2 was identified as S phase cells, which are distinguished by an increase in Geminin-Clover expression, reaching medium levels. Population 3 consisted of cells in G2 phase, which exhibited the highest expression of Geminin-Clover. Population 4 consisted of cells undergoing the transition to mitosis, which was evidenced by the progressive loss of SLBP-Turquoise2 expression. Furthermore, we identified a fifth population that was characterized by high levels of SLBP-Turquoise2 and Cdt1-mKO2 that had not been previously characterized with the FUCCI system. The intensity of Ki67 in the nucleus increased with cell cycle progression. In contrast, p21 levels were significantly low in proliferating cells, reaching a minimum in S phase. In addition, the intensity of H3 was found to double from S phase onward, consistent with the DNA duplication process (Figure 1B). We observed that the fifth population, newly identified in this study, had the lowest Ki67 levels (Figure 1B-D), strong p21 expression (Figure 1B and E-F), and lower intensity of H3 similar to cells in G1 (Figure 1B and G-H). Based on these characteristics, we concluded that this population had exited the cell cycle and was henceforth referred to as the G0. However, the observation of a lower intensity of H3 in G0 cells was unexpected, as it could be indicative of a loss of genomic material.

**Figure 1.**
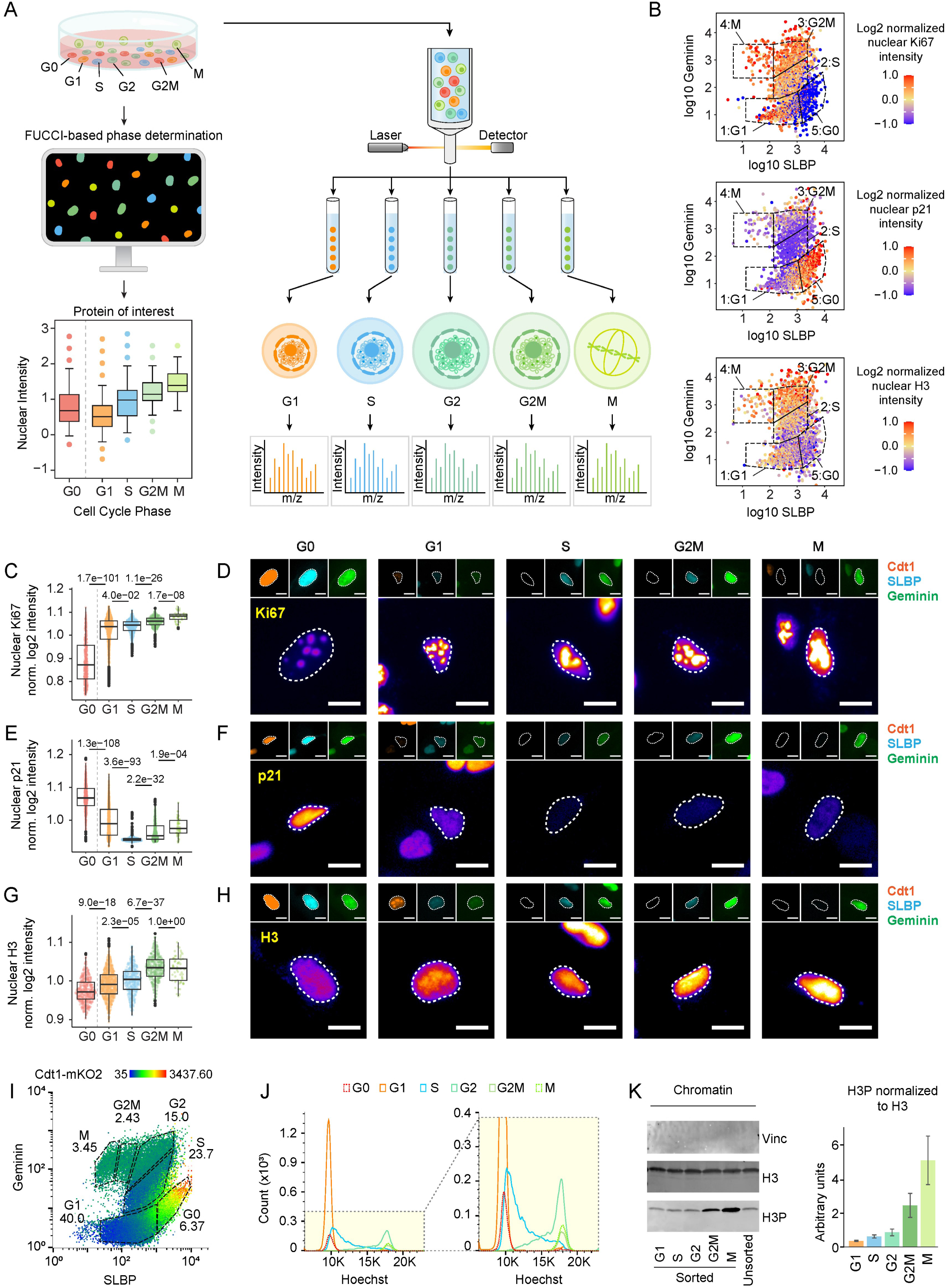
Characterization of U2OS-FUCCI-3 reporter system for the analysis of cell cycle-dependent chromatin associations. (A) Experimental overview: U2OS-FUCCI3 cells were cultured to maintain a stable distribution across different cell cycle phases. Subsequently, they were either subjected to microscopy for immunofluorescence or live-cell imaging studies or sorted by FACS to obtain protein extracts for mass spectrometry analysis. (B) Representation of Ki67, p21 and H3 nuclear intensity measured by immunofluorescence and plotted accordingly to FUCCI-3-determined cell cycle phases. (C) Immunofluorescence-based quantification of the intensity of Ki67 in the nuclear region identified as the integration of signal from the FUCCI-3 fluorescent markers, which has been used for the determination of cell cycle phases. 3 biological replicates were analyzed (*n*G0 = 476, *n*G1 = 890, *n*S = 572, *n*G2/M = 370, *n*M = 50; outliers removed, 3 SD; unpaired two-tailed Wilcoxon test). (D) Representative images showing the changes of Ki67 inside the nucleus during cell cycle progression, as well as the respective signal of the FUCCI-3 fluorescent markers of the illustrated cell, which confirm the belonging to the assigned cell cycle phase. (E) Immunofluorescence-based quantification of the intensity of p21 in the nuclear region identified as the integration of signal from the FUCCI-3 fluorescent markers, which has been used for the determination of cell cycle phases. 3 biological replicates were analyzed (*n*G0 = 559, *n*G1 = 768, *n*S = 384, *n*G2/M = 424, *n*M = 55; outliers removed, 3 SD; unpaired two-tailed Wilcoxon test). (F) Representative images showing the changes of p21 inside the nucleus during cell cycle progression, as well as the respective signal of the FUCCI-3 fluorescent markers of the illustrated cell, which confirm the belonging to the assigned cell cycle phase. (G) Immunofluorescence-based quantification of the intensity of H3 in the nuclear region identified as the integration of signal from the FUCCI-3 fluorescent markers, which has been used for the determination of cell cycle phases. 3 biological replicates were analyzed (*n*G0 = 490, *n*G1 = 912, *n*S = 468, *n*G2/M = 439, *n*M = 58; outliers removed, 3 SD; unpaired two-tailed Wilcoxon test).(H) Representative images showing the changes of H3 inside the nucleus during cell cycle progression, as well as the respective signal of the FUCCI-3 fluorescent markers of the illustrated cell, which confirm the belonging to the assigned cell cycle phase. (I) Comparison between FUCCI-3 based identification of cell cycle phases and (J) FACS-based cell cycle profiles obtained with Hoechst staining. (K) Western blot validation of the chromatin fractionation using Vinculin as cytoplasmic marker and H3 as chromatin marker accompanied by the and quantification of Phosphorylated H3 (H3P, serine 10) normalized to H3 during cell cycle.

Given that the choice of microscopy over FACS also enabled the analysis of nuclear characteristics, we incorporated the size of the nucleus into the evaluation of the expression of these cell cycle markers. As expected, cell cycle progression was accompanied by an increase in nuclear size (Figure S1D). The evaluation of nuclear size allowed us to calculate the total amount of a protein present in the nucleus at different phases of the cell cycle (referred to as integrated intensity, calculated as mean intensity multiplied by nuclear size). This is different from protein concentration, which is determined by the mean intensity of the protein of interest. We observed that the behavior of Ki67 and p21 during the cell cycle was similar when considering the mean intensity (Figure 1C-F) or the integrated intensity (Figure S1E). In proliferating cells, H3 concentration and protein amounts also showed similar behavior. However, while the H3 concentration decreased in G0 cells compared to G1 cells (Figure 1G-H), the total amount of H3 protein was the same in G0 and G1 cells (Figure S1E). This effect can be explained by the enlarged nuclei that characterize the G0 population (Figure S1C) and rules out the possibility that G0 cells lost genetic material. We further validated the state of the U2OS FUCCI-3 reporter cell line by comparing the integration of the fluorescent markers with Hoechst staining followed by FACS analysis. As expected, the DNA content matched with each of the FUCCI-3 determined cell cycle phases. Moreover, the FUCCI-3 reporter enabled discrimination of G2, G2/M and M phases that could not be achieved with the Hoechst staining alone (Figure 1I-J). In contrast to Hoechst which is a DNA intercalating agent, the FUCCI-3 system does not impair cell viability over time, thereby enabling the tracking of cells without affecting the cell cycle. To test the suitability of the FUCCI-3 reporter cell line for cell cycle tracking, we used drugs that arrest the cell cycle at specific phases, to analyze the downstream cell cycle defects. Cells were treated with either RO-3306, a cyclin-dependent kinase 1 (CDK1) inhibitor, which arrests cells at the G2/M boundary^35^, and the tubulin polymerization inhibitor nocodazole ^36^, which blocks cells in prometaphase. By performing live cell imaging, we observed that cell proliferation was affected by both treatments (Figure S1F). Moreover, tracking the different reporter expressions allowed us to identify the cell cycle arrest at the expected phases (Figure S1G-H), confirming the suitability of the generated reporter for live cell imaging studies. Finally, we tested the suitability of the FUCCI-3 reporter system for viable FACS sorting. Five populations corresponding to the cell cycle phases and transitions of proliferating cells (G1, S, G2, G2/M, and M) were sorted according to the expression of FUCCI-3 fluorescent markers (Figure S1I). Following sorting, each population was released into culture and monitored via live cell imaging for a period of 40 hours to confirm cell viability and correct cell cycle progression (Figure S1J). Finally, an in-house-optimized chromatin purification protocol was performed on each of the sorted populations, followed by sonication and endonuclease digestion with benzonase to extract the chromatin-associated proteins, which we refer to as the chromatome^26,31,27^ (Figure S1K). To adapt the protocol for the sorted M population, it was necessary to avoid the nuclear lysis step, given that mitotic cells lack a nuclear membrane. The chromatin pellet was obtained after cellular membrane lysis, washed and sonicated prior benzonase digestion. The purity of the protein samples and their assignment to specific cell cycle phases was validated by Western blot using Vinculin as a cytoplasmic marker, H3 as a chromatin marker, and phosphorylated H3 (pH3, serin 10) as a G2-M marker (Figure 1K and Figure S1L).

In summary, these results showed that the optimized FUCCI-3 reporter system allowed the generation of a stable cell line expressing cell cycle reporters for FACS and imaging-based analysis that can be used for phenotypic and molecular dissection of cell cycle related events.

### Chromatome-mass spectrometry across cell cycle progression

To identify cell cycle-dependent changes in the chromatome, data-independent acquisition mass spectrometry (DIA-MS) was performed on the sorted populations. Following normalization of the raw data (Figure S2A) principal component analysis (PCA) was performed and demonstrated a robust clustering of biological replicates, which exhibited notable differences between sorted populations and from the unsorted and unsynchronized cells (Figure S2B). Furthermore, a relative compartment enrichment analysis was conducted to validate the purity of the chromatome fractions (Figure S2C). Thereafter, we performed a cluster analysis of proteins showing cell cycle oscillatory behavior, selecting those proteins that showed a significant change between at least two consecutive phases (Figure S2D-E). For instance, cluster number 1 grouped proteins increasing from G1 to M phase, with a pronounced increase between G1 and S phase. Gene ontology (GO) analysis of this cluster enriched for terms related to general cell cycle progression and regulation (Figure S2F). On the other hand, cluster number 11 grouped proteins that showed an increase towards the end of the cell cycle and enriched for GO terms related to chromosome segregation (Figure S2F). These results validated our experimental setup for high-throughput identification of cell cycle-dependent chromatin-bound proteins. To understand whether nuclear metabolism could be related to cell cycle progression, we performed a metabolism-centric analysis, searching our cell cycle chromatome dataset for metabolic genes and regulators as either members of the generic genome-scale metabolic model of Homo sapiens^37^ or targets of the human CRISPR-Cas9 metabolic library^38^ (Figure 2A, Figure S3A). We identified several metabolism-related factors that change on chromatin during the cell cycle, including metabolic regulators and canonical metabolic enzymes. Among metabolic regulators we retrieved Aurora kinases (AURKA and AURKB) that use ATP as phosphate group donor^39,40^, thus impinging on cellular metabolism. Moreover, Aurora A has been directly linked to metabolic processes via interaction with the ATP synthases^41^ or the phosphorylation of Pyruvate Kinase M2 (PKM2)^42^. Aurora Kinases showed minimal chromatin levels in G1 and progressively increased to reach their maximum in G2/M (Figure 2B, Cluster 5; Figure S3A, Cluster 3 violet). Moreover, epigenetic factors involved in chromatin methylation balance, including DNA (cytosine-5)-methyltransferase 1 (DNMT1), Histone lysine methyltransferase 5A (KMT5A), Histone lysine methyltransferase Suppressor Of Variegation 3-9 Homolog 1 (SUV39H1), and Lysine demethylase 5B (KDM5B), demonstrated dynamic fluctuations on chromatin during the cell cycle (Figure 2B and Figure S3A). Given their impact on the equilibrium of available methyl groups, these enzymes exert a profound influence on metabolic processes^43^, thereby classifying them as metabolic regulators. SUV39H1 showed a linear increase during cell cycle (Figure S3A, violet), similar to AURKA/B. In contrast, DNMT1, KMT5A and KDM5B exhibited an increase at specific cell cycle phases (Figure 2B and Figure S3A; DNMT1 pine green, KMT5A and KDM5B in light green). DNMT1 appeared to increase its concentration on chromatin from the G1 phase to the S phase, subsequently returning to baseline levels from the G2/M phase onwards. In contrast, KMT5A and KDM5B exhibited an inverse pattern, reaching a minimal level in S phase and increasing from G2 onward. We thus sought to determine whether the observed behavior was related to a differential binding of these proteins to chromatin during the cell cycle, a variation of their absolute nuclear abundance, or to oscillations of their nuclear concentration across cell cycle progression. To address this question, we validated our DIA-MS results with high-throughput immunofluorescence staining of U2OS FUCCI-3 cells using antibodies specific for either DNMT1, KMT5A, or KDM5B. The fixed and stained populations exhibited normal FUCCI-3 cell cycle profiles (Figure S3B-D). We observed that the nuclear integrated intensity of each protein, which measures the absolute amount of protein present in the nuclear compartment, increased across the cell cycle, following the proportional increase in the size of the nucleus (Figure S3E-G). It is noteworthy that the total amount of DNMT1 decreased in the G0 population in the nucleus, indicating a decrease in the availability of DNMT1 for the same amount of DNA (Figure S3E). This finding may be consistent with the reduced DNA methylation often observed in senescent cells^44,45,46,47,48^, suggesting that the population we called G0 in the FUCCI-3 plot may be composed of both quiescent and senescent cells, as both express high levels of p21^49,50,51^. Conversely, we observed a marked increase of KMT5A (Figure S3F), and KDM5B in G0 cells (Figure S3G). The differences detected in protein abundance between G0 and G1 cells were also observed when considering protein concentration, suggesting that the decrease or increase was independent of the size of the nucleus (Figure 2C-E). In cycling cells, however, the size of the nucleus made a difference. Indeed, DNMT1 levels increased from G1 to S and then stabilized, whereas KMT5A and KDM5B levels only increased significantly from G2/M (Figure 2D-E). This may suggest that the size of the nucleus partially controls the nuclear, and possibly chromatin, concentration of these epigenetic factors. In agreement with this hypothesis, the integration of the chromatome DIA-MS (Figure 2B) and high-throughput immunofluorescence analysis (Figure 2C-E) indicated that DNMT1 exhibited an increase from the G1 to S phase in both the nuclear compartment (high-throughput immunofluorescence) and the chromatin compartment (chromatome DIA-MS). In addition, while the intensity of the nuclear protein signal remained stable during progression to G2/M phase, there was a decrease in chromatin binding, indicating that DNMT1 chromatin extrusion occurred progressively as chromosomes condensed towards mitosis. In the case of KMT5A and KDM5B, on the other hand, we observed a correlation between the decreased chromatin binding in the S phase and the lowest nuclear concentration observed in high-throughput immunofluorescence. Similarly, chromatin binding and nuclear concentration increased in G2, exhibiting levels comparable to those observed in G1, and reached maximum levels in M (Figure 2B and Figure 2C-E).

**Figure 2.**
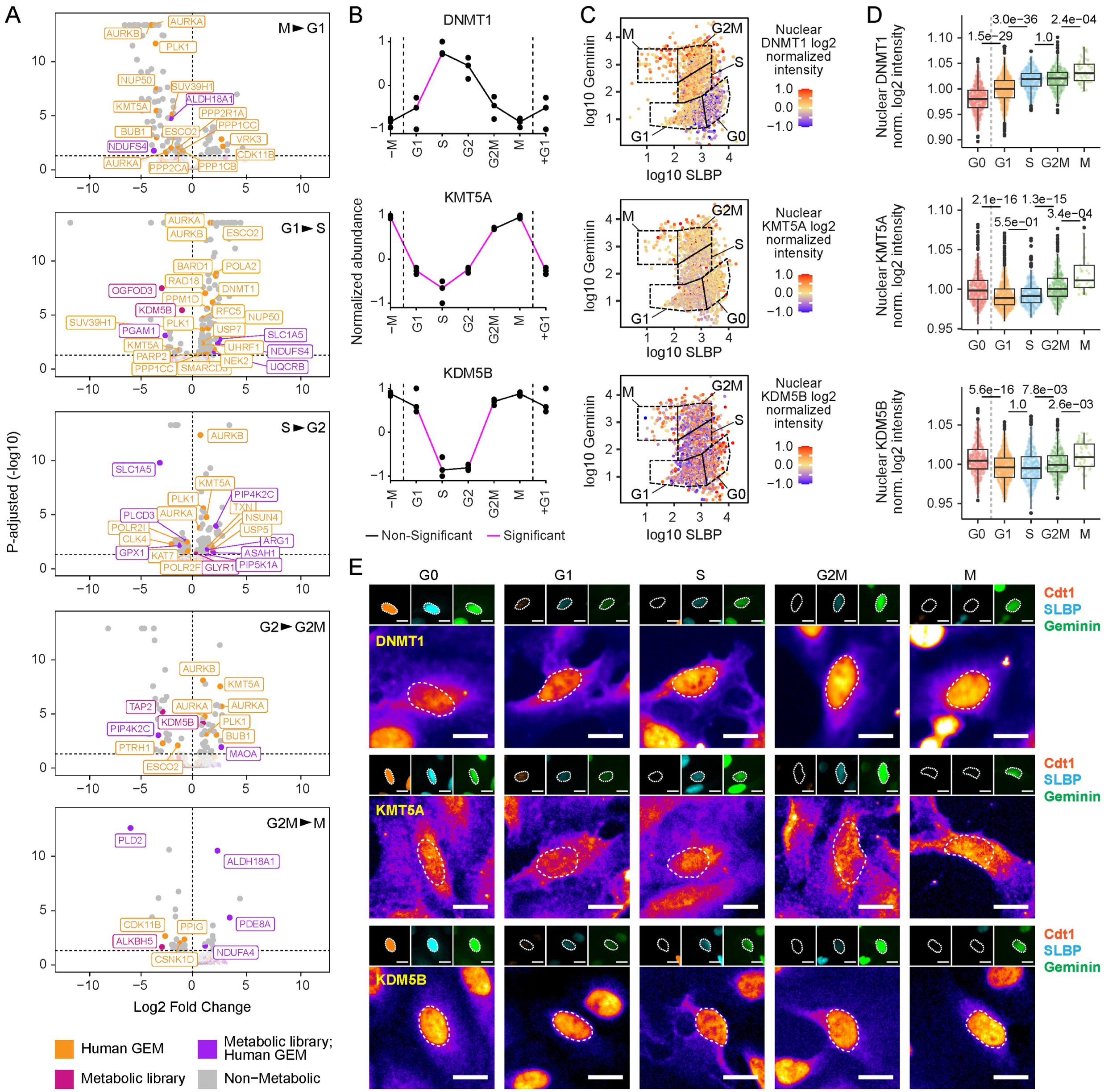
Cell cycle control of chromatin and nuclear localization of methylation-related enzymes. (A) Volcano plots of mass spectrometry data showing the chromatin enrichment and depletion of proteins related to metabolism. Proteins belonging to the human genome-scale metabolic model (GEM)^37^ are represented in orange, proteins belonging to the metabolic sgRNA library^38^ are represented in pink, proteins belonging to both groups are represented in violet. In gray are proteins not related to metabolism. (B) Relative abundance of DNMT1, KMT5A and KDM5B on chromatin across cell cycle as determined by mass spectrometry on chromatin fractions. Magenta lines mean that the difference between the 2 consecutive phases was significant. Black lines mean that no significant difference was found between consecutive phases. Statistical analysis was performed according to the test_diff function in DEP R package^112^, following intensity imputation, which internally uses limma statistical testing^125^. (C) Representation of DNMT1, KMT5A or KDM5B nuclear intensity measured by immunofluorescence and plotted accordingly to FUCCI-3-determined cell cycle phases. (D) Immunofluorescence-based quantification of nuclear DNMT1, KMT5A or KDM5B intensities across cell cycle phases determined with the FUCCI-3 system. 3 biological replicates were analyzed (DNMT1 IF: *n*G0 = 510, *n*G1 = 772, *n*S = 423, *n*G2/M = 348, *n*M = 55; KMT5A IF: *n*G0 = 463, *n*G1 = 675, *n*S = 375, *n*G2/M = 374, *n*M = 35; KDM5B IF: *n*G0 = 436, *n*G1 = 720, *n*S = 412, *n*G2/M = 315, *n*M = 47; outliers removed, 3 SD; unpaired two-tailed Wilcoxon test). (E) Representative images showing the changes of DNMT1, KMT5A or KDM5B inside the nucleus during cell cycle progression, as well as the respective signal of the FUCCI-3 fluorescent markers of the illustrated cell, which confirm the belonging to the assigned cell cycle phase.

Using our workflow, which discriminates between proteins enriched in the nuclear compartment and those bound to chromatin during different phases of the cell cycle, we were able to unravel the cell cycle-associated dynamics of epigenetic enzymes involved in DNA and histone methylation. Moreover, we suggested that changes in nuclear size during cell cycle may control the nuclear concentration of these enzymes and thus their availability on chromatin.

### Phosphatidylinositol metabolism associates with chromatin in a cell cycle-dependent manner

The metabolism of phosphatidylinositol is strictly connected to availability of methylation substrates^52,53^ and is well documented for its implications in cell signaling and the dynamics of cellular membranes, including those of the plasma membrane, endosomes, lysosomes, and the Golgi apparatus^54,55^. Phosphatidylinositol phosphates and related enzymes have been demonstrated to localize in the nucleus^56,57^, although their nuclear role remains understudied. For example, the nuclear localization of phosphatidylinositol 4,5-bisphosphate (PIP2) is known to change during the cell cycle^58,59^. However, the reason for this cell cycle-dependent behavior of nuclear PIP2 is not yet understood. The results of our chromatome DIA-MS analysis demonstrated the presence of a significant number of PIP2-related enzymes on chromatin during the cell cycle. Furthermore, the nuclear presence of these enzymes appears to outflow into the regulation of histone methylation (Figure 3A). To confirm the presence of nuclear and cell cycle-dependent PIP2 metabolism, we characterized the cell cycle-associated behavior of phosphatidylinositol 4-phosphate 5-kinase type 1 alpha (PIP5K1A), which phosphorylates phosphatidylinositol 4-phosphate (PI4P) to PIP2^60,61^ and is thus a central component of PIP2 metabolism. Chromatome DIA-MS analysis showed that PIP5K1A was bound to chromatin mainly in the G1 and G2 phase, while its chromatin association decreased during the S and after the G2 phase (Figure 3A-B and Figure S3A, pine green). Nevertheless, high-throughput immunofluorescence using a PIP5K1A-specific antibody on U2OS FUCCI-3 cells, exhibiting a typical cell cycle distribution (Figure S4A), revealed that the nuclear concentration of PIP5K1A remained stable from G1 to S phase and increased in G2/M (Figure 3C-D and Figure S4B). On the other hand, the nuclear abundance of the protein increased in a cell cycle-dependent manner (Figure S4C). The different behavior of the chromatin and nuclear pools indicated that while the nuclear concentration of the protein is kept stable from G1 to S, its chromatin association is cell cycle regulated. To confirm this, we sorted U2OS FUCCI-3 cells into either G1, S, or G2, isolated the cytosol, nucleoplasm, and chromatin fractions, and performed Western blot analysis. The results confirmed that while nucleoplasm concentration was stable in G1 and S and increased in G2, chromatin association was much stronger in G1 and G2 than in S (Figure 3E). This suggests that PIP5K1A is reloaded onto chromatin after DNA duplication. We also performed high-throughput immunofluorescence of the metabolic enzymes phospholipase C delta 3 (PLCD3) and phospholipase D2 (PLD2), which use PIP2 either as a substrate or as a modulator downstream of PIP5K1A, respectively. PLCD3 catalyzes the hydrolysis of PIP2 to produce the second messengers diacylglycerol (DAG) and inositol 1,4,5-trisphosphate (IP3). DAG activates protein kinase C (PKC), a well-known cell cycle regulator^62^. IP3 triggers the release of intracellular calcium^63^, which regulates the progression from G1 to S phase of the cell cycle^64^ and the activity of calmodulin-dependent kinases, including some cyclin-dependent kinases (CDKs)^65,64^. In addition, IP3 receptor activity has recently been shown to regulate spindle orientation, implicating that IP3 is important for proper mitotic progression^66^. On the other hand, PIP2 binding to PLD2 enhances its activity in the hydrolysis of phosphatidylcholine to phosphatidic acid (PA) and choline^67^. Choline participates in methionine synthesis via betaine homocysteine methyltransferase (BHMT), thereby regulating levels of the methyl donor S-adenosylmethionine (SAM) and influencing histone and DNA methylation^68^. In addition, PA is required for the *de novo* synthesis of cell membranes and cell growth^69^, which are essential for the cell cycle. As they compete for PIP2, increased PLCD3 activity inhibits PLD2^67^. In addition, the synthesis of PA by PLD2 stimulates the activity of PIP5K1A to produce PIP2^67^ (Figure 3A). Our chromatome DIA-MS data showed that PLCD3 is predominantly found on chromatin at the beginning of the cell cycle and during M phase (Figure 3A-B and Figure S3A, yellow). These findings are consistent with a role for DAG and IP3 in regulating G1 to S progression and for IP3 in regulating spindle orientation. However, when we performed high-throughput immunofluorescence in U2OS FUCCI-3 cells displaying a normal cell cycle profile (Figure S4D), we observed that the nuclear concentration of PLCD3 remained stable from G1 to S while increasing in G2 (Figure 3F-G and Figure S4E). On the other hand, its abundance increased throughout the cell cycle (Figure S4F), following an opposite behavior to that observed for the chromatin-bound pool. In addition, the PLD2 chromatin-bound fraction decreased in the M phase (Figure 3A-B and Figure S3A, pine green), whereas high-throughput immunofluorescence on normal cycling U2OS FUCCI-3 cells (Figure S4G) showed that PLCD2 nuclear concentration was stable from G1 to S, but increased in G2 and M (Figure 3H-I and Figure S4H). On the other hand, the nuclear abundance of the protein increased in a cell cycle-dependent manner (Figure S4I). The different nuclear and chromatin behavior of PLCD3 and PLD2, coupled with the fact that they regulate each other’s activity by competing for PIP2, suggests that their cell cycle-associated roles may be regulated not only by their nuclear localization but also by their respective subnuclear compartmentalization. Given the differential presence of PIP2-related enzymes in the chromatin and nuclear environment during cell cycle progression, we quantified the level of PIP2 itself in the nucleus by integrating the quantification of the FUCCI signal (Figure S4J) with a PIP2-specific antibody (Figure S4K). Our data showed that nuclear PIP2 concentration was largely stable from G1 to G2/M and increased in M (Figure 3J-K and Figure S4K), whereas nuclear PIP2 abundance progressively increased during the cell cycle (Figure S4L). Notably, PIP2-related enzymes and PIP2 itself showed higher nuclear concentrations in G0 phase than in G1 phase (Figure 3 C-K). This indicates that cell cycle arrest potentially correlates with an accumulation of PIP2 metabolism in the nuclear milieu, which may be linked to an imbalance in chromatin states.

**Figure 3.**
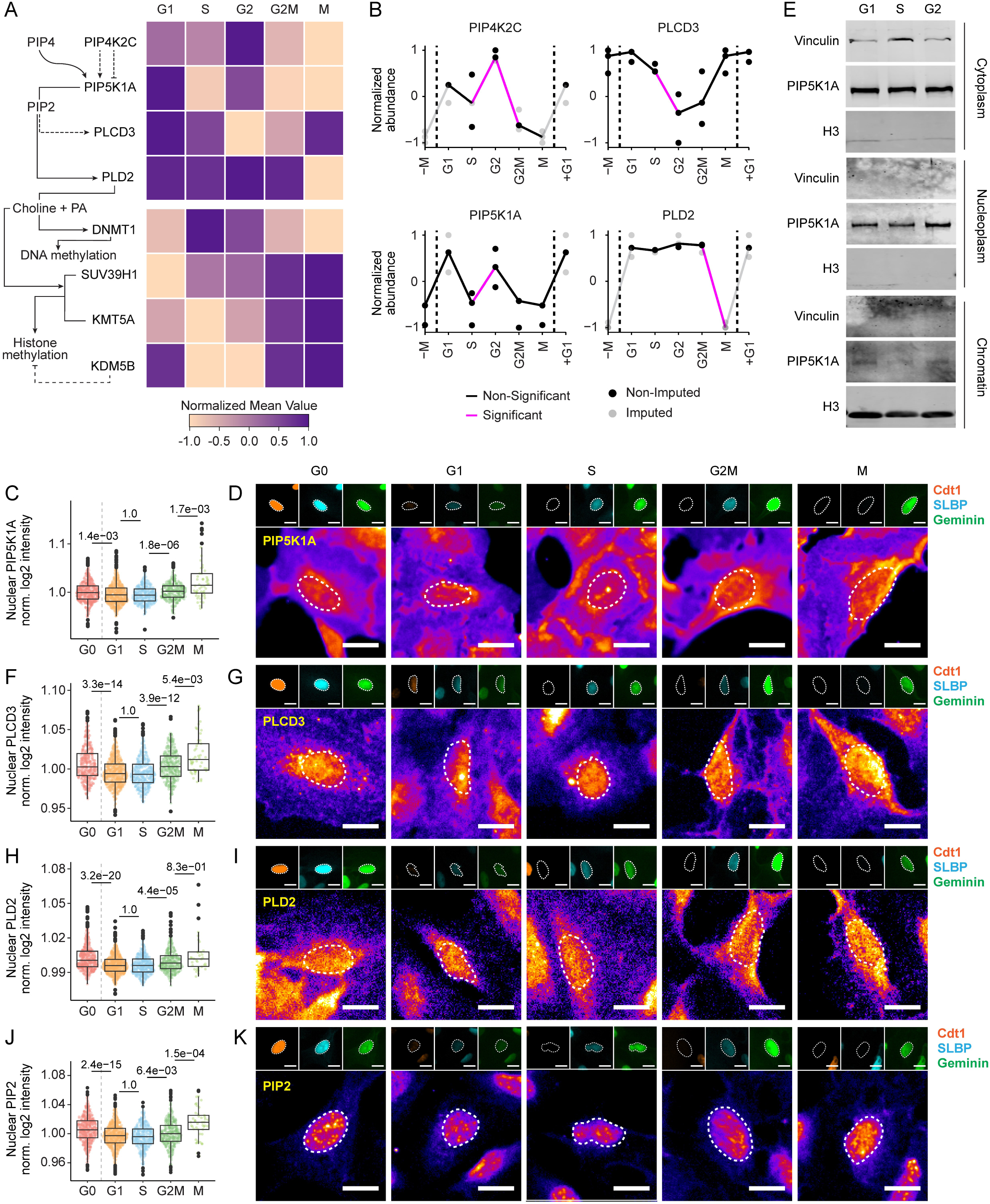
Phosphatidylinositol metabolism oscillates on chromatin during the cell cycle. (A) Reconstruction of a chromatin-associated metabolic network representing the integration of phosphatidylinositol metabolism and chromatin methylation, obtained by plotting the normalized mean value of the abundance of each protein determined by mass spectrometry in each cell cycle phase. (B) Relative chromatin abundance of PIP5K1A, PLCD3 and PLD2 across cell cycle as determined by mass spectrometry on chromatin fractions. Magenta lines mean that the difference between the 2 consecutive phases was significant. Black lines mean that no significant difference was found between consecutive phases. Semi-transparent dots correspond to imputed values. Statistical analysis was performed according to the test_diff function in DEP R package^112^, following intensity imputation, which internally uses limma statistical testing^125^. (C) Immunofluorescence-based quantification of nuclear PIP5K1A intensities across cell cycle phases determined with the FUCCI-3 system. 3 biological replicates were analyzed (*n*G0 = 530, *n*G1 = 633, *n*S = 380, *n*G2/M = 334, *n*M = 52; outliers removed, 3 SD; unpaired two-tailed Wilcoxon test). (D) Representative images showing the changes of PIP5K1A inside the nucleus during cell cycle progression, as well as the respective signal of the FUCCI-3 fluorescent markers of the illustrated cell, which confirm the belonging to the assigned cell cycle phase. (E) Western blot showing enrichment of PIP5K1A in the chromatin, nuclear and cytoplasmic fractions at different cell cycle stages. H3 and Vinculin are used as chromatin and cytoplasmic controls, respectively. (F) Immunofluorescence-based quantification of nuclear PLCD3 intensities across cell cycle phases determined with the FUCCI-3 system. 3 biological replicates were analyzed (*n*G0 = 438, *n*G1 = 685, *n*S = 444, *n*G2/M = 367, *n*M = 49; outliers removed, 3 SD; unpaired two-tailed Wilcoxon test). (G) Representative images showing the changes of PLCD3 inside the nucleus during cell cycle progression, as well as the respective signal of the FUCCI-3 fluorescent markers of the illustrated cell, which confirm the belonging to the assigned cell cycle phase. (H) Immunofluorescence-based quantification of nuclear PLD2 intensities across cell cycle phases determined with the FUCCI-3 system. 3 biological replicates were analyzed (*n*G0 = 479, *n*G1 = 758, *n*S = 421, *n*G2/M = 342, *n*M = 34; outliers removed, 3 SD; unpaired two-tailed Wilcoxon test). (I) Representative images showing the changes of PLD2 inside the nucleus during cell cycle progression, as well as the respective signal of the FUCCI-3 fluorescent markers of the illustrated cell, which confirm the belonging to the assigned cell cycle phase. (J) Immunofluorescence-based quantification of nuclear PIP2 intensities across cell cycle phases determined with the FUCCI-3 system. 3 biological replicates were analyzed (*n*G0 = 514, *n*G1 = 724, *n*S = 406, *n*G2/M = 347, *n*M = 45; outliers removed, 3 SD; unpaired two-tailed Wilcoxon test). (K) Representative images showing the changes of PIP2 inside the nucleus during cell cycle progression, as well as the respective signal of the FUCCI-3 fluorescent markers of the illustrated cell, which confirm the belonging to the assigned cell cycle phase.

Our results indicate that the enzymes involved in PIP2 metabolism in the nucleus and on chromatin are differentially regulated during the cell cycle, suggesting that they may have specific functions dependent on their subnuclear localization. In addition, enzymes involved in PIP2 metabolism and PIP2 itself increase in the nucleus in G0, suggesting that an imbalance of nuclear PIP2 may correlate with cell cycle arrest.

### Inhibition of phosphatidylinositol metabolism compromises cell cycle and impairs nucleoli dynamics

In the nucleus, PIP2 has been found to differentially localize in Fibrillarin areas and speckles^70,71,72,73,74^, with accumulation in Fibrillarin areas characterized by small and more concentrated PIP2 foci, whereas in speckles by larger foci with less signal intensity, which we could confirm by immunofluorescence (Figure S5A). Fibrillarin is a core component of the nucleolus and is closely linked to cell cycle regulation^75^. Consistent with this, Fibrillarin has been associated with proper S-phase progression^76^. In line with the observation that nucleolar activity increases as the cell cycle progresses, we noted that Fibrillarin regions themselves (used as a proxy for nucleoli) exhibited expansion in size throughout the cell cycle, similar to nuclear regions that did not contain Fibrillarin (Figure S5B). However, Fibrillarin concentration in nucleoli remained unaltered (Figure S5C-D), indicating that maintaining a stable concentration of Fibrillarin within nucleoli is crucial during cell cycle progression. Therefore, we aimed at investigating changes of PIP2 nuclear localization during cell cycle progression. To do so, we integrated high-throughput immunofluorescence-based DNA quantification (via Hoechst staining) and nuclear dimension to determine the Hoechst integrated intensity, and use it to identify the cell cycle phases of the analyzed cells (Figure S5E), as previously shown^71,70^. This method was applied to U2OS wild-type cells, which allowed staining with two different fluorescent secondary antibodies, in contrast to U2OS FUCCI-3 cells, which allowed visualization of only one protein at a time. The disadvantage of this method is that it does not allow distinguishing cells at the boundary between two phases or in mitosis, as is possible using the FUCCI-3 system (Figure 1J). In addition, cells cannot be tracked in live cell imaging. By performing PIP2 immunofluorescence in combination with Fibrillarin staining, we observed that the nuclear distribution of PIP2 changed during the cell cycle. While the area of each PIP2 foci did not considerably change either inside or outside the nucleoli (identified by Fibrillarin staining), the number of PIP2 foci in nucleoli increased from G1 to S and decreased from S to G2, whereas there was no change outside the nucleoli (Figure 4A). This suggests that cell cycle progression alters PIP2 distribution in a subnuclear-specific manner. Integration of these data with the cell cycle associated chromatin levels of PIP5K1A, the enzyme that synthesizes PIP2, (Figure 3A) suggested that PIP2 synthesis on chromatin occurs mainly during G1 and G2, while it is stored in nucleoli in the S phase. Consequently, we targeted PIP2 metabolism by inhibiting PLCD3 and PIP5K1A to see how this would impact on cell cycle progression. The FUCCI-3 cells were treated with U-73122 (PLC inhibitor) or ISA-2011B (PIP5K1A inhibitor) and the respective IC50s were determined over a period of 72 hours in order to select a concentration that would not induce extensive cell death for the following experiments (Figure S5F). The administration of near-IC50 concentrations of both compounds to FUCCI-3 U2OS cells for 60 hours resulted in a strong reduction in cell proliferation (Figure 4B and Figure S5G). Thus, we sought to ascertain the differential impact of the inhibitions on the cell cycle progression by live cell imaging. Inhibition of PLCD3 resulted in an initial accumulation of cells in the G2/M phase, which persisted for up to 12 hours, followed by a progressive arrest in the G1/S phase after 48 hours (Figure 4C). This is in accordance with the double PLCD3 peak observed with chromatome DIA-MS, which demonstrated the presence of this enzyme in the G1, S and M phases (Figure 3A-B). Conversely, the inhibition of PIP5K1A resulted in the immediate accumulation of cells in G1 (Figure 4C), the phase in which the highest abundance of this enzyme on chromatin was observed (Figure 3A-B). The results indicated that the presence of these enzymes on chromatin at precise cell cycle phases could be a prerequisite for the correct progression of the cell cycle via PIP2 synthesis and consumption. When treating the cells for 48 hours with U-73122 or ISA-2011B, we observed that the effect on cell proliferation was moderate in comparison to 60 or 72 hours of treatment (Figure S5H). Accordingly, this specific time point was selected for the analysis of early alterations in nuclear PIP2 metabolism. PLCD3 inhibition did not result in any alterations to PLCD3 levels within the nucleus (Figure S5I-K). Conversely, PIP5K1A inhibition led to a slight yet consistent increase in nuclear PIP5K1A levels (Figure S5L-N). This phenomenon could potentially represent a cellular mechanism employed to counteract the effects of the inhibitor within the nucleus. Since PIP5K1A is directly involved in PIP2 synthesis, we aimed to analyze potential alterations in the subnuclear distribution of PIP2 in response to PIP5K1A inhibition. In U2OS wild-type cells we observed that the concentration of nuclear PIP2 did not change overall upon PIP5K1A inhibition (Figure 4D). However, we noticed a drastic rearrangement of the nucleolar area, which increased significantly, as did Fibrillarin intensity, while the number of nucleoli decreased (Figure 4E-H). This indicated that nucleoli underwent fusion and subsequent enlargement as a consequence of the treatment. The analysis of the PIP2 signal inside nucleoli revealed a reduction in the concentration of PIP2 in these Fibrillarin-positive regions in G1 and G2/M (Figure 4I). This finding is consistent with the evidence that PIP5K1A is more abundant on chromatin during G1 and G2, while it is excluded from chromatin during S phase (Figure 3A and E), and suggests that PIP5K1A chromatin localization may control the amount of PIP2 specifically in nucleoli in a cell cycle-dependent manner.

**Figure 4.**
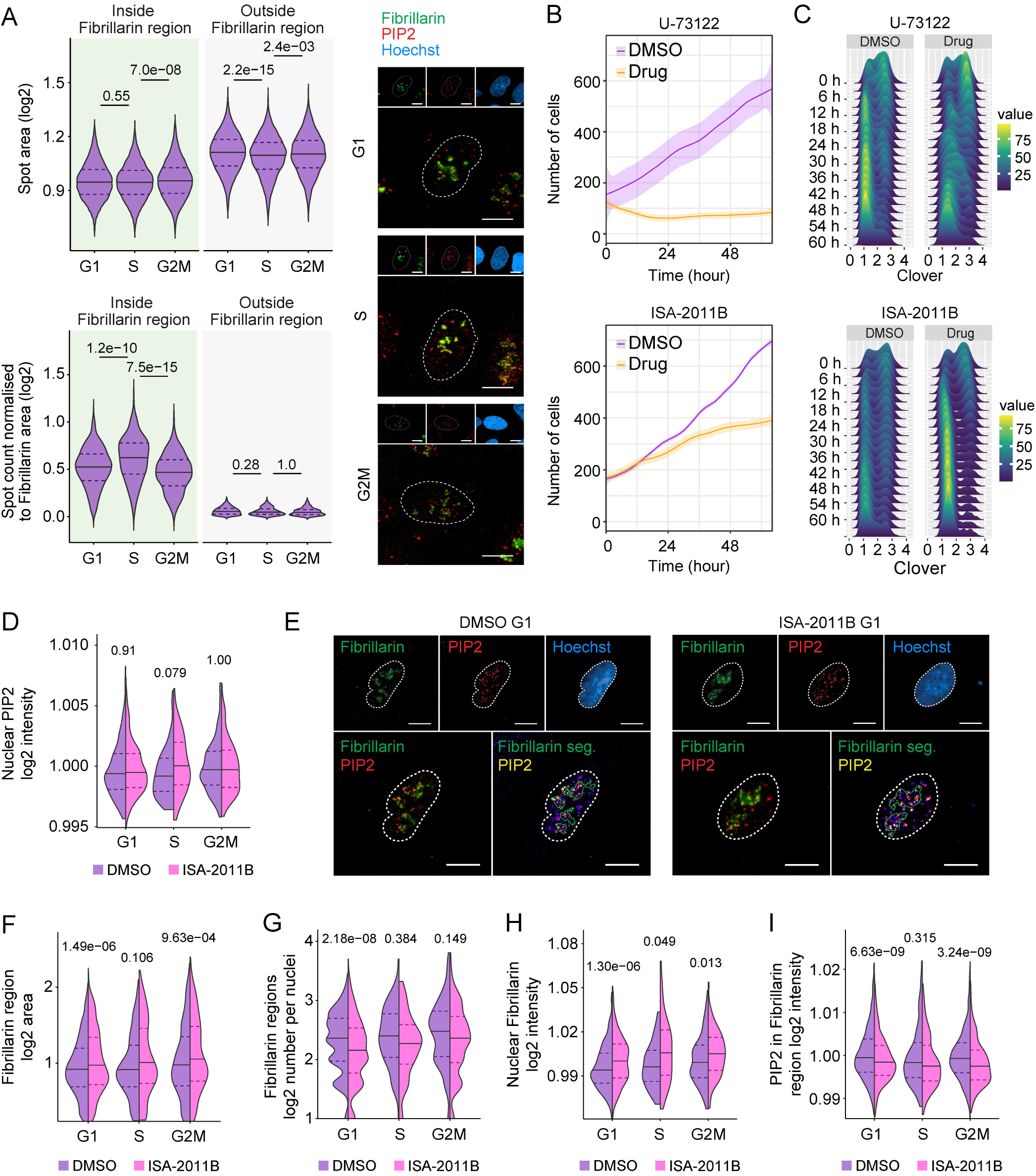
PIP2 nucleolar localization is dependent on nuclear phosphatidylinositol metabolism and cell cycle regulation. (A) Immunofluorescence-based quantification of the area of PIP2 foci (top panel) and their number normalized to the Fibrillarin area size (bottom panel), classified as either inside nucleoli (identified by Fibrillarin staining) or outside nucleoli (Fibrillarin-negative regions). Experiments were performed in U2OS cells grouped by cell cycle phase using Hoechst integrated intensity. 3 biological replicates were analyzed (*n*G1 = 1704, *n*S = 678, *n*G2M = 489; outliers removed, 3 SD; unpaired two-tailed Wilcoxon test). The panels on the right are representative images showing the intensity and distribution of Fibrillarin (green) and PIP2 (red) in each identified cell cycle phase. The nucleus, identified with Hoechst staining, is shown in blue. (B) Growth curve of U2OS cells treated with DMSO (negative control), U-73122 (PLCD3 inhibitor; 5.54 μM) or ISA-2011B (PIP5K1A inhibitor; 41.33 μM). FUCCI-3 U2OS cells were treated for 60 hours and nuclei were identified by the expression of FUCCI-3 fluorescent markers and counted over time to determine cell growth. Tracking was performed with a minimum of 150 cells per treatment. (C) Live cell imaging of cells treated with U-73122 (5.54 μM) or ISA-2011B (41.33 μM) and monitored for 60 hours. Changes in the number of Clover-Geminin negative (G1) and positive (S & G2) cells are shown in the ridge plot, illustrating the cell cycle distribution from the beginning of the treatment and every 6 hours. (D) Immunofluorescence-based quantification of nuclear PIP2. (E) Representative images showing the intensity and distribution of Fibrillarin (green) and PIP2 (red) in the G1 phase in cells treated with DMSO or ISA-2011B (41.33 μM). The nucleus, identified with Hoechst staining, is shown in blue. The segmentation of the Fibrillarin region is shown as a green outline in the merged image, while PIP2 is shown in FIRE_lut. Scale bar is 10µm. (F) Immunofluorescence-based quantification of nucleoli area (Fibrillarin segmented region), (G) number of nucleoli per nucleus, (H) intensity of Fibrillarin in nuclei, (I) intensity of PIP2 in nucleoli. The above quantifications (E-I) were performed in cell cycle stratified population based on Hoechst integrated intensity after DMSO or ISA-2011B (41.33 μM) treatments. 3 biological replicates were analyzed and the minimum number of cells used for *n*DMSO_G1 = 568, *n*ISA-2011B_G1 = 471, *n*DMSO_S = 77, *n*ISA-2011B_S = 49, *n*DMSO_G2M = 188 and *n*ISA-2011B_G2M = 144 (outliers removed, 3 SD; unpaired two-tailed Wilcoxon test).

These findings demonstrate that disrupting the metabolism of nuclear PIP2 exerts a profound influence on the regulation of the cell cycle. Furthermore, the inhibition of PIP5K1A, which synthesizes PIP2, results in a specific reduction of PIP2 concentration in the nucleolar regions.

### A functional link between nuclear PIP2 metabolism and chromatin methylation

Nucleoli are integral to the distribution and function of heterochromatin in the nucleus, serving as hubs that facilitate chromatin organization and silencing^77,78^. The enlargement of these Fibrillarin regions, together with the differential behavior of PIP2 inside and outside nucleoli upon inhibition of PIP5K1A, suggested a possible effect on chromatin organization upon treatment. To test this hypothesis, we analyzed the monomethylation of Histone 4 lysine 20 (H4K20me1), which is deposited by KMT5A^79^, whose chromatin association is regulated by the cell cycle as illustrated before (Figure 3A-B). H4K20me1 is deposited at the centromeric region during mitosis and allows the recruitment of centromeric and kinetochore proteins^79,80^. In addition, we tested the constitutive heterochromatin mark Histone 3 lysine 9 trimethylation (H3K9me3), which is deposited by SUV39H1 that progressively increases on chromatin from S phase to mitosis (Figure 3A-B). Finally, we included the facultative heterochromatin methylation mark Histone 3 lysine 27 trimethylation (H3K27me3), which is deposited by the Enhancer Of Zeste 1/2 Polycomb Repressive Complex 2 subunits (EZH1 and EZH2), which were not found in our database to be chromatin-bound in a cell cycle regulated manner. To impinge on PIP2 metabolism, we treated U2OS wild-type cells with PLCD3 and PIP5K1A inhibitors and measured the nuclear amount of the mentioned histone marks to identify differential behaviors in a cell cycle dependent manner. Firstly, we observed that H4K20me1 exhibited a progressive increase from the S phase onwards, reaching an expected maximum in the M phase (Figure 5A). Following the inhibition of PLCD3 for a period of 48 hours (Figure S6A), the levels of H4K20me1 exhibited an increase across all phases, with the exception of M (Figure 5A and Figure S6B). This data indicated that inhibiting PLCD3 activity may result in the accumulation of PIP2, which in turn stimulates PLD2 and enhances choline synthesis. Choline acts as a methyl donor in the synthesis of methionine, thereby contributing to the synthesis of SAM^68^. This suggests that the maintenance of H4K20me1 from G1 to G2/M may depend on PIP2 and choline availability. To test this hypothesis, we inhibited PIP2 synthesis for 48 hours via PIP5K1A inhibition (Figure S6C). The treatment had the opposite effect on H4K20me1 compared to PLCD3 inhibition, revealing a decrease in nuclear levels in all cell cycle phases except for the M phase (Figure 5B and Figure S6D). This supported the hypothesis that increasing PIP2 availability may be a requirement to balance SAM levels during the cell cycle via choline-derived methionine synthesis. When analyzing the levels of H3K9me3 in untreated cells, we observed that they were stable from the G1 to G2/M phase and increased in the M phase, probably due to chromatin compaction that leads to chromosome condensation. Inhibition of PLCD3 for 48h (Figure S6E), did not lead to a significant change in the levels of H3K9me3 (Figure 5C, S6F). PIP5K1A inhibition for 48h (Figure S6G) significantly increased the H3K9me3 mark in every phase except the M phase (Figure 5D, S6H), in contrast to the effect observed for H4K20me1. This may indicate that the synthesis of PIP2 and choline is not a prerequisite for the maintenance of constitutive heterochromatin, which may rely more on folate metabolism or the methionine salvage pathway for SAM synthesis. However, the observed increased levels of constitutive heterochromatin following the inhibition of PIP2 synthesis suggested that PIP2 metabolism is important for maintaining the correct level of chromatin compaction throughout the cell cycle. Similar to the observations made for H3K9me3, levels of H3K27me3 remained unchanged from the G1 to G2/M phase and showed an increase in the M phase in untreated conditions. However, we found that inhibition of PLCD3 or PIP5K1A for 48h (Figure S6I-J), both resulted in increased levels of H3K27me3 from the G1 to G2/M phase, in contrast to what was seen for H3K9me3 levels (Figure 5E-F and Figure S6K-L). These results suggest that choline is also not essential for the maintenance of this facultative heterochromatin mark, which as constitutive heterochromatin may rely more on folate metabolism or the methionine salvage pathway for SAM synthesis. However, disruption of the PIP2 balance led to an increase in facultative heterochromatin, again suggesting that PIP2 metabolism is essential for maintaining a balance in chromatin compaction during the cell cycle.

**Figure 5.**
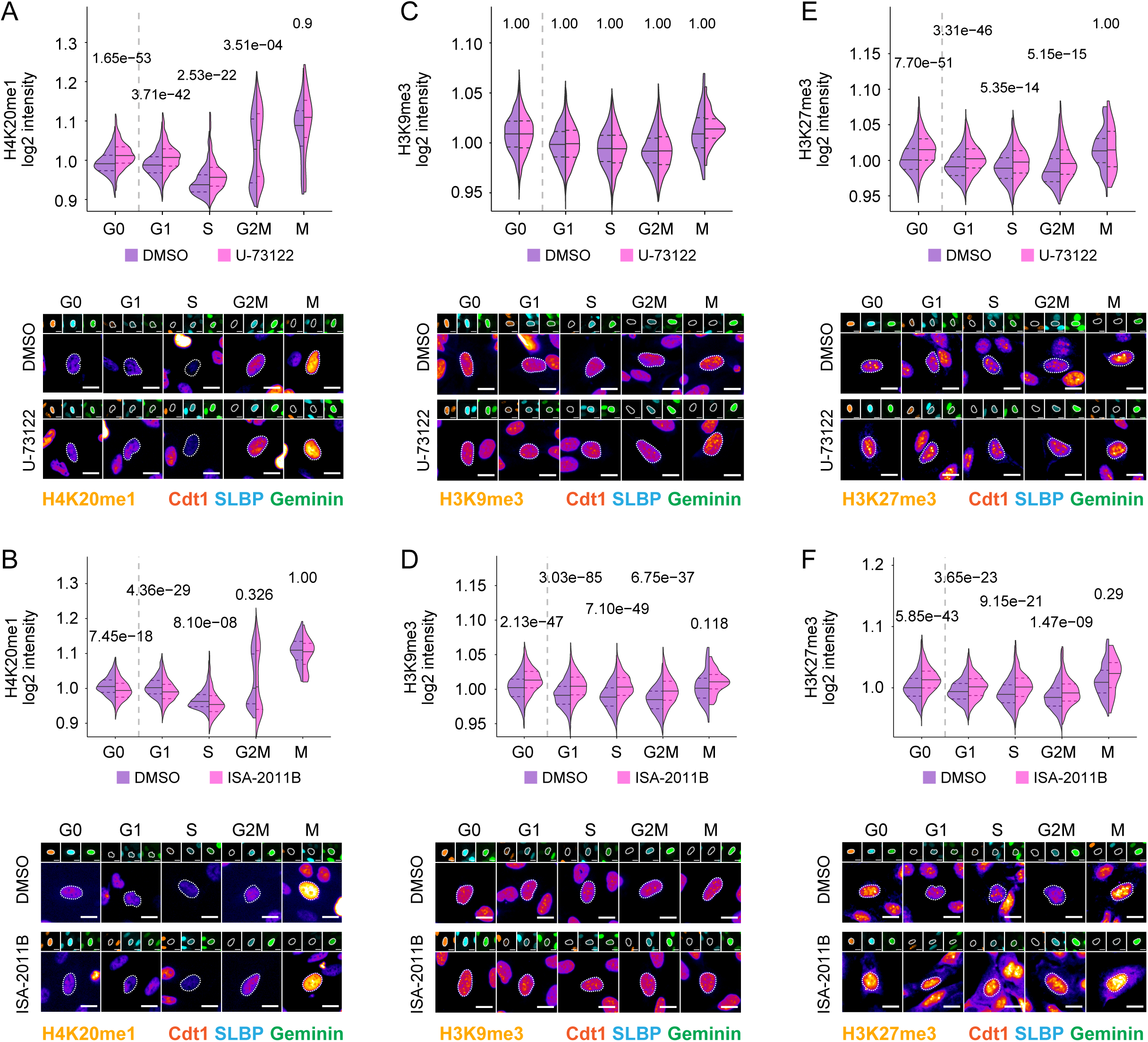
PIP2 perturbations differentially affect histone methylation. (A) Immunofluorescence-based quantification of H4K20me1 intensities across cell cycle phases determined with the FUCCI-3 system in cells treated with DMSO or U-73122 (5.54 μM). 3 biological replicates were analyzed (*n*DMSO_G0 = 1045, *n*U-73122_G0 = 1376, *n*DMSO_G1 = 1100, *n*U-73122_G1 = 1274, *n*DMSO_S = 708, *n*U-73122_S = 811, *n*DMSO_G2M = 613, *n*U-73122_G2M = 628, *n*DMSO_M = 38, *n*U-73122_M = 29; outliers removed, 3 SD; unpaired two-tailed Wilcoxon test). The lower panels display representative images showing the changes of H4K20me1 in the nucleus during cell cycle progression, as well as the corresponding signal of the FUCCI-3 fluorescent marker of the depicted cell, confirming the assigned cell cycle phase. (B) Immunofluorescence-based quantification of H4K20me1 intensities across cell cycle phases determined with the FUCCI-3 system in cells treated with DMSO or ISA-2011B (41.33 μM). 3 biological replicates were analyzed (*n*DMSO_G0 = 878, *n*ISA-2011B_G0 = 961, *n*DMSO_G1 = 1644, *n*ISA-2011B_G1 = 1205, *n*DMSO_S = 933, *n*ISA-2011B = 423, *n*DMSO_G2M = 823, *n*ISA-2011B_G2M = 469, *n*DMSO_M = 72, ISA-2011B_M = 27; outliers removed, 3 SD; unpaired two-tailed Wilcoxon test). The lower panels display representative images showing the changes of H4K20me1 in the nucleus during cell cycle progression, as well as the corresponding signal of the FUCCI-3 fluorescent marker of the depicted cell, confirming the assigned cell cycle phase. (C) Immunofluorescence-based quantification of H3K9me3 intensities across cell cycle phases determined with the FUCCI-3 system in cells treated with DMSO or U-73122 (5.54 μM). 3 biological replicates were analyzed (*n*DMSO_G0 = 1310, *n*U-73122_G0 = 1421, *n*DMSO_G1 = 2087, *n*U-73122_G1 = 1359, *n*DMSO_S = 1100, *n*U-73122 = 919, *n*DMSO_G2M = 721, *n*U-73122_G2M = 671, *n*DMSO_M = 64, U-73122_M = 25; outliers removed, 3 SD; unpaired two-tailed Wilcoxon test). The lower panels display representative images showing the changes of H3K9me3 in the nucleus during cell cycle progression, as well as the corresponding signal of the FUCCI-3 fluorescent marker of the depicted cell, confirming the assigned cell cycle phase. (D) Immunofluorescence-based quantification of H3K9me3 intensities across cell cycle phases determined with the FUCCI-3 system in cells treated with DMSO or ISA-2011B (41.33 μM). 3 biological replicates were analyzed (*n*DMSO_G0 = 933, *n*ISA-2011B_G0 = 1474, *n*DMSO_G1 = 1686, *n*ISA-2011B_G1 = 1573, *n*DMSO_S = 1001, *n*ISA-2011B = 671, *n*DMSO_G2M = 857, *n*ISA-2011B_G2M = 6720, *n*DMSO_M = 91, ISA-2011B_M = 39; outliers removed, 3 SD; unpaired two-tailed Wilcoxon test). The lower panels display representative images showing the changes of H3K9me3 in the nucleus during cell cycle progression, as well as the corresponding signal of the FUCCI-3 fluorescent marker of the depicted cell, confirming the assigned cell cycle phase. (E) Immunofluorescence-based quantification of H3K27me3 intensities across cell cycle phases determined with the FUCCI-3 system in cells treated with DMSO or U-73122 (5.54 μM). 3 biological replicates were analyzed (*n*DMSO_G0 = 1298, U-73122_G0 = 1198, *n*DMSO_G1 = 1763, *n*U-73122_G1 = 1134, *n*DMSO_S = 970, *n*U-73122 = 724, *n*DMSO_G2M = 741, *n*U-73122_G2M = 512, *n*DMSO_M = 53, U-73122_M = 27; outliers removed, 3 SD; unpaired two-tailed Wilcoxon test). The lower panels display representative images showing the changes of H3K27me3 in the nucleus during cell cycle progression, as well as the corresponding signal of the FUCCI-3 fluorescent marker of the depicted cell, confirming the assigned cell cycle phase. (F) Immunofluorescence-based quantification of H3K27me3 intensities across cell cycle phases determined with the FUCCI-3 system in cells treated with DMSO or ISA-2011B (41.33 μM). 3 biological replicates were analyzed (*n*DMSO_G0 = 1032, *n*ISA-2011B_G0 = 1308, *n*DMSO_G1 = 1877, *n*ISA-2011B_G1 = 1435, *n*DMSO_S = 1052, *n*ISA-2011B = 630, *n*DMSO_G2M = 945, *n*ISA-2011B_G2M = 5168, *n*DMSO_M = 116, ISA-2011B_M = 28; outliers removed, 3 SD; unpaired two-tailed Wilcoxon test). The lower panels display representative images showing the changes of H3K27me3 in the nucleus during cell cycle progression, as well as the corresponding signal of the FUCCI-3 fluorescent marker of the depicted cell, confirming the assigned cell cycle phase.

In conclusion, the data presented indicate that there is a relationship between the structure of nucleoli, the nuclear metabolism of PIP2 and histone methylation during the cell cycle. Moreover, distinct methionine and SAM pools appear to be susceptible to differential modulation by PIP2 metabolism, with the maintenance of specific heterochromatin types contingent upon the nature of the pool in question.

## Discussion

Recent studies in transcriptomics and proteomics have investigated cell cycle oscillations and demonstrated that gene expression and protein synthesis change during the cell cycle^81,82,83,84,85^. However, despite the importance of protein compartmentalization in the regulation of protein functions^86^, protein compartmentalization in the context of cell cycle regulation has not been studied. Given the recent importance of nuclear compartmentalization of metabolic enzymes in chromatin functions^23,24,25,26,27,28,29,30,31,87^, here we asked whether metabolic enzymes could be relocated either on chromatin or in the nucleus in a cell cycle-dependent manner. By integrating high-throughput immunofluorescence and mass spectrometry, we have established an unbiased approach that allows the discrimination of subpools of nuclear metabolic enzymes that are either in the nucleoplasm or on chromatin in different cell cycle phases. Moreover, by focusing on PIP2 metabolism as a showcase, we proposed the possibility that subnuclear accumulation of PIP2 may influence cell cycle progression through global changes in histone methylation.

In this study, we established a system based on the generation of stable FUCCI reporters that allows the tracking of cells during the cell cycle over generations without impairing cell proliferation (Figure 1). Unlike other studies^88,89,90,91,92,93^, our FUCCI-3 system allowed naive sorting of cells in different cell cycle phases and cell cycle phase transitions without the need to treat them with cell cycle inhibitors, which could cause severe artifacts due to the cell cycle arrest phenotype they induce. With the sorted populations, we performed quantitative mass spectrometry on the chromatin fraction (the chromatome) to look for factors that oscillate during the cell cycle. Since it is not possible to accurately estimate how cell volume and its protein content vary during the cell cycle, our mass spectrometry analysis was performed considering equal amounts of proteins. However, given that the absolute amount of protein tends to increase as the cell cycle progresses^84^, it is important to consider that the measured oscillations reflect the relative amount of a target protein compared to the total protein content and do not necessarily reflect changes in the absolute amount of the target.

We found that the nuclear and chromatin localization of chromatin methylation regulators (Figure 2) and phosphatidylinositol pathway enzymes (Figure 3) are interrelated and regulated by the cell cycle. Our workflow integrated nuclear imaging with chromatome data to identify fluctuations between the nuclear and chromatin pools of such factors. This detailed analysis revealed a functional link between cell cycle progression, phosphatidylinositol synthesis and histone methylation (Figure 3 and Figure 4). Finally, in the last part of the study, we analyzed the behavior of Fibrillarin in the nucleus during the cell cycle as a proxy for nucleoli dynamics. Since PIP2 colocalizes with Fibrillarin, we integrated the changes of PIP2 abundance in the nucleoli with the changes in histone methylation, thus drawing a connection between subnuclear PIP2 metabolism and chromatin dynamics throughout the cell cycle (Figure 5).

We were able to detect cell cycle arrested cells using the FUCCI-3 system (Figure 1). Interestingly, nuclear PLCD3 and PLD2, as well as PIP2 itself, were increased in cells exhibiting a cell cycle arrested phenotype (Figure 3C-K and Figure S5J), suggesting that nuclear accumulation of PIP2 and PIP2-consuming metabolic enzymes may be dysfunctional and affect cell fitness. Enlarged Fibrillar regions, also observed in this study by blocking PIP2 synthesis with inhibition of PIP5K1A (Figure 4H), have previously been implicated in aging and senescence^94,95^. Taken together, these data may link nucleolar integrity to nuclear PIP2 metabolism, identifying this functional connection as a feature of actively proliferating cells. Furthermore, the relationship with PIP2 and histone methylation may suggest that altered subnuclear PIP2 metabolism or storage may be involved in promoting cell cycle progression or arrest, providing another way in which dysregulation of nuclear PIP2 metabolism may affect the cell cycle. Moreover, although not discussed in this study, the fact that Fibrillarin is a ribosomal RNA methyltransferase^96,97^ raises the question of whether the imbalance of nucleolar PIP2 could affect ribosomal RNA methylation and whether this could be related to cell cycle arrest.

Our study not only links protein compartmentalization to cell cycle progression, but also raises the possibility that subnuclear pools of metabolic enzymes and metabolites may be associated with cell states and fates. In this context, it would be interesting to investigate in the future whether differential nuclear PIP2 localization might be present in cells that exit the cell cycle and undergo differentiation, or whether cancer cells are able to reactivate subnuclear PIP2 metabolism that allows cell cycle reentry in an uncontrolled manner. Beyond PIP2 metabolism, spatial metabolomics and proteomics may in the future help to elucidate how proteins and metabolites are compartmentalized during the cell cycle, once the technical limitations that currently prevent the quantification of subnuclear pools of enzymes or their metabolites have been overcome.

## Materials and Methods

### Cloning of FUCCI-3 lentiviral vectors

To generate the FUCCI-3 lentiviral plasmids, the neomycin and hygromycin coding regions driven by the SV40 promoter were PCR amplified from the pcDNA3^98^ and GW209_pCRIS-PITChv2-C-d vectors^99^, respectively, with primers 1 and 2, defined below, using Phusion High-Fidelity DNA Polymerase (ThermoFisher Scientific; #F530). The resulting products were purified using the QIAquick Gel Extraction Kit (Qiagen, 28706). The PCR products and the backbones pLL3.7m-mTurquoise2-SLBP and pLL3.7m-Clover-Geminin(1-110)-IRES-mKO2-Cdt were cut with MluI-HF (NEB, R3198S), followed by dephosphorylation with rSAP (NEB, M0371S). The cut vectors were gel purified using the QIAquick Gel Extraction Kit (Qiagen, 28706). The PCR products were cloned into each lentiviral vector overnight by classical ligation with T4 ligase (NEB, M0202S). The resulting ligation product was transformed in 25 µl of DH5α (Thermo Fisher Scientific, 18265017) and selected at 100 µg/ml in house ampicillin plates. Individual colonies were analyzed by Sanger sequencing using primers 3, 4 and 5, defined below. The coding sequence of Histone 1 fused to Maroon1 from pLL3.7m-mTurquoise2-SLBP was eliminated by excision of the vector with EcoRI-HF (NEB, R3101S). After gel purification using the QIAquick Gel Extraction Kit (Qiagen, 28706), the resulting cut vector was religated overnight with T4 ligase (NEB, M0202S). The resulting ligation product was transformed with 25 µl of DH5α (Thermo Fisher Scientific, 18265017) and selected at 100 µg/ml in house ampicillin plates. Individual colonies were analyzed by Sanger sequencing using primer 6, defined below.

Integrated DNA technologies oligos:

Forward sequence Primer 1

5’-CAATGACTTACAAGGCAGCTGACGCGTGTGTGTCAGTTAGGGTGTGGAAAG-3’

Reverse sequence Primer 2

5’-CTTTTCTTTTAAAATCACGCGTTAAGATACATTGATGAGTTTGGACAAACCA-3’

Forward sequence Primer 3

5’-ATCGGCACTTTGCATCGGC -3’

Forward sequence Primer 4

5’-GCAGGGTCGATGCGACG -3’

Reverse sequence Primer 5

5’-CAGCAGCTACCAATGCTGATT -3’

Reverse sequence Primer 6

5’-CTAGGTACAATTCGATATCAAGCTTATCG -3’

### FUCCI-3 reporter cell line generation

To establish a stable FUCCI-3 system, cells were infected with lentiviral particles containing the modified plasmids derived from the original FUCCI-4 system^100^: pLL3.7m-mTurquoise2-SLBP(18-126)-Neomycin and pLL3.7m-Clover-Geminin(1-110)-IRES-mKO2-Cdt(30-120)-Hygromycin. Following transduction, cells were cultured in media containing 200 µg/mL Geneticin (Thermo Scientific; #10131027) and 150 µg/mL Hygromycin (Sigma-Aldrich; #H3274) to select for successfully transduced cells, with the selection process lasting four to seven days. Fluorescence-activated cell sorting (FACS) using a BD Influx sorter was performed to enrich cells displaying proper FUCCI-3 expression dynamics. This FUCCI-3 system was specifically designed to track the cell cycle by monitoring three cell cycle-regulated proteins: Clover-Geminin, SLBP-Turquoise2, and Cdt1-mKO2. To ensure the long-term correct expression of the FUCCI-3 reporter system and homogeneity of the population, the FUCCI-3 reporter cells were not maintained in culture for more than one month. Another batch of cells was then thawed and sorted for the expected color pattern of the FUCCI-3 reporter system.

### Cell culture

The U2OS cell line, either wild-type or transduced with the FUCCI-3 reporter system, was cultured in Petri dishes at 37°C and 5% CO2 with DMEM (Gibco; #11966025) supplemented with 10% Fetal Bovine Serum (FBS) (Gibco; #10270106) and 1% penicillin/streptomycin (Gibco; #15140122).

### Lentiviral particles production and cell transduction

Lentiviral particles were generated using HEK293T cells cultured in 150 mm dishes and transfected via a polyethyleneimine (PEI)-mediated protocol (Polysciences; #23966-1). Briefly, 5.5 µg of the pCMV-dR8_91 plasmid and 4.2 µg of the pVSV-G envelope plasmid were combined with 8.4 µg of the plasmid carrying the gene of interest in 1 mL of Opti-MEM medium (Gibco; #11058021). Separately, 54.6 µL of 1 mg/mL PEI solution was diluted with 900 µL of Opti-MEM. After 5 minutes, the plasmid mix and the PEI solution were combined, incubated for 20 minutes to form transfection complexes, and added dropwise to the HEK293T cells in DMEM (Gibco; #11966025) supplemented with 10% Fetal Bovine Serum (FBS, Gibco; #10270106) and 1% penicillin/streptomycin (Gibco; #15140122). Six hours later, the medium was replaced with fresh growth medium. Supernatant containing lentiviral particles was harvested 48 hours post-transfection. For lentiviral transduction of U2OS cells, 1 mL of viral supernatant was added per well of a 6-well plate, along with Polybrene (Sigma-Aldrich; #TR-1003) at a final concentration of 10 µg/mL.

### FACS sorting

To ensure proper expression of the FUCCI-3 fluorescent markers in the U2OS-FUCCI3 cell line or to collect cells according to their cell cycle phase, the cells were incubated for at least 48 hours after the final splitting to obtain a homogeneous and asynchronous population. Subsequently, the cells were treated with Trypsin (Thermo Scientific; #25200072), resuspended in DMEM (Gibco; #11966025) supplemented with 10% Fetal Bovine Serum (FBS, Gibco; #10270106) and 1% penicillin/streptomycin (Gibco; #15140122), and placed on ice. The cells were sorted according to their cell cycle stage, as indicated by the FUCCI-3 fluorescent markers, on a BD Influx Cell Sorter (BD Biosciences) with a stable temperature of 4°C. The 457 nm laser and CFP parameter were used for SLBP-Turquoise2 detection, the 488 nm laser and FITC parameter were used for Clover-Geminin detection, and the 561 nm laser and PE parameter were used for Cdt1-mKO2 detection. For the FUCCI-3 system’s validation, a 355 nm laser was employed to detect Hoechst and the results were analyzed with Flowjo_V10.

### Immunofluorescence

Immunofluorescence experiments were conducted by seeding cells on transparent, flat-bottom 96-or 384-well plates (Revvity; #6055302, #6057302) and incubating them for a minimum of 48 hours prior to fixation. The fixation was conducted by the addition of 16% methanol-free formaldehyde (Thermo Fisher Scientific; #28908) to the culture, resulting in a final concentration of 4%, for a period of 10 minutes. The cells were then washed three times with PBS (1x) and permeabilized with 0.2% Triton X-100 in PBS (1x) for 20 minutes. The preparation was blocked with 5% bovine serum albumin (BSA) diluted in PBS (1x) for a period of 40 minutes. Subsequently, the cells were incubated with the primary antibodies for a period of 2.5 hours, after which they were washed once with 0.05% Tween 20 in PBS (1x) and twice with PBS (1x). The secondary antibodies were then applied for a period of 45 minutes, after which the cells were washed once with 0.05% Tween 20 in PBS (1x) and twice with PBS (1x). Furthermore, for U2OS wild-type cells, a 10-minute incubation with Hoechst 33342 (Thermo Fisher Scientific; #H3570) diluted in PBS (1x) was conducted for DNA staining.

The following primary antibodies were used: Ki67 (Cell Signaling; #9129; 1:400), p21 (BD Bioscience; #AB_396415; 1:200), H3 (Cell Signaling; #14269), DNMT1 (ProteinTech; #24206-1-AP; 1:100), KMT5A (Thermo Fisher Scientific; #PA5-31467; 1:250), KDM5B (Cell Signaling; #3273; 1:250), PIP5K1A (ProteinTech; #15713-1-AP; 1:200), PLCD3 (ProteinTech; #16792-1-AP; 1:250), PLD2 (Cell Signaling; #13904; 1:250), Fibrillarin (Cell Signaling; #2639; 1:400), PIP2 (Thermo Fisher Scientific; #MA3-500; 1:200), H4K20me1 (Diagenode; #C15410034; 1:500), H3K9me3 (Diagenode; #C15410193; 1:1000), and H3K27me3 (Diagenode; #C15410195; 1:200). The following secondary antibodies were used: Alexa Fluor 488 donkey anti-rabbit (Thermo Fisher Scientific; #A21206; 1:1000), Alexa Fluor 647 goat anti-rabbit (Thermo Fisher Scientific; #A21244; 1:1000) and Alexa Fluor 633 goat anti-mouse (Thermo Fisher Scientific; #A21050; 1:1000).

### High-Throughput Microscopy

Immunofluorescent images were captured using the Operetta High Content Screening System (PerkinElmer) with a 10X, 20X, or 63X objective in either non-confocal or confocal mode. Live-cell images were acquired with the 20x objective, maintaining a temperature of 37°C and 5% CO_2_. Images were quantified using the Harmony software (version 4.9) and the results analyzed with R programming environment (v 4.4).

#### Cell cycle phase determination

To determine the cell cycle phase based on the FUCCI-3 system, the excitation filter 390-420 and emission 430-500 was used to measure SLBP-Turquoise2 signal, the excitation filter 460-490 and emission 500-550 was used to measure Clover-Geminin signal, and the excitation filter 530-560 and emission 570-650 was used to measure Cdt1-mKO2 signal. The signal of the 3 colors was merged using Harmony’s building block *Calculate Image* and the nuclei were segmented based on this image using the building block *Find Nuclei*. The nuclei crossing the edge of the image were discarded with *Select Population* and the option *Remove Borders Objects*, and their roundness and area were determined with *Calculate Morphology Properties*. The incorrect or low-quality segmented nuclei were visually inspected and removed by applying thresholds taking into account the roundness and size in the *Select Population* building block. A ring region of 8 pixels was further determined around the segmented nuclei for measuring the background using the building block *Select Cell Region*. The mean intensity for each of the 3 FUCCI fluorescent markers, in the nuclei and their corresponding ring region, was measured with the building block *Calculate Intensity Properties*. The results were exported in a tab separated table, which was used as input for a custom R script to perform further calculations including the compensation for the bleed through signal between Clover and Turquoise2 and subtraction of the background signal (signal from the ring region) from the corresponding nuclei. A scatterplot, named FUCCI-3 plot, was then generated with the *ggplot2*^101^ package by representing the nuclear mean intensity signal of the 3 FUCCI fluorescent markers: log10 Clover-Geminin in the Y axis, the log10 SLBP-Turquoise2 in the X axis and the log2 normalized Cdt1-mKO2 as 3 color scale of the dots. To ensure the visualization of all the dots representing a nucleus, the *geom_voronoi* option from the *ggforce*^102^ package was used. Based on the FUCCI-3 plot pattern, cell cycle gates were drawn. Across different experiments, the shape of the overall pattern varied depending on the specific acquisition conditions. To keep the gating criteria as homogeneous as possible between experiments, the data and the gates were further transformed to show a similar pattern in the control conditions (untreated, DMSO controls or 0h timepoint). The coordinates from each gate’s vertex were used in the *points_in_polygon* tool from the *sp* package^103,104^ to identify in which gate each nucleus was contained, and this was added as a new feature (Phase_Group column) for each nuclei with the *mutate* tool from the *dplyr*^105^ package.

To determine the cell cycle phase based on Hoechst staining, we used a strategy adapted from previously published methods^106,107^. Briefly, the excitation filter 355-385 and emission 430-500 was used to measure Hoechst signal, and the nuclei were segmented based on this image using the building block *Find Nuclei*. The nuclei crossing the edge of the image were discarded with *Select Population* and the option *Remove Borders Objects*, and their roundness and area were determined with *Calculate Morphology Properties*. The incorrect or low-quality segmented nuclei were visually inspected and removed by applying thresholds taking into account the roundness and size in the *Select Population* building block. A ring region of 8 pixels was further determined around the segmented nuclei for measuring the background using the building block *Select Cell Region*. The results were exported in a tab separated table, which was used as input for a custom R script to perform further calculations. A histogram was then generated with the *ggplot2*^101^ package by representing the nuclear mean intensity signal in the X axis. The histogram was visually inspected to find the signature peaks of G1 and G2 typical and thresholds were determined for the Hoechst signal; these were then used to classify the nuclei into the different cell cycle phases with the *mutate* tool from the *dplyr*^105^ package.

#### Quantification of nuclear proteins

The mean intensity, interpreted as the concentration of the targeted protein in the overall nucleus, was calculated using the building block *Calculate Intensity Properties* and selecting the filtered segmented nuclei as population. The integrated intensity, interpreted as the total amount of protein in the nuclei, was calculated from multiplying the mean intensity times the area of the nucleus in R.

#### Data Normalization

To represent the oscillation of protein intensity during the cell cycle in "FUCCI plots," raw measurements were normalized by dividing the intensity of each nucleus by the population mean, followed by a log2 transformation. The color scale was adjusted to range from -1 to 1, emphasizing populations with low and high intensities. For boxplots, violin plots, and statistical comparisons across treatments and/or cell cycle phases, normalization was performed to account for biological replicates. This was done by dividing each object’s intensity value by the population mean (in log2 scale), ensuring data consistency and enabling reliable comparisons across biological replicates.

#### Data Representation

To represent any feature of interest in the context of cell cycle FUCCI plots were generated. As described before, these plots consist of representing log10 Clover-Geminin in the Y axis, the log10 SLBP-Turquoise2 in the X axis and the log2 normalized Cdt1-mKO2 as 3 color scale of the dots. However, once the cell cycle phases have been visually determined, the color dimension is used for the representation of the feature of interest. These plots were generated with the *ggplot2* package^101^ and the *geom_vorono*i option from the *ggforce* package^102^. Boxplots were generated using *geom_boxplot* and each dot representing a nucleus was generated with *geom_sina* from the *ggforce* package^102^. Violin plots were generated using the functions *geom_split_violin*, *create_quantile_segment_frame* and *GeomSplitviolin* (original function by Jan Gleixner (@jan-glx), adjustments by Wouter van der Bijl (@Axeman)). Final aesthetic adjustments were done with Illustrator CS5 (Adobe).

#### Region segmentation and analysis of foci and colocalization

PIP2 spots were segmented using Harmony’s building block *Find Spots*, using as population the filtered segmented nuclei and restricting the analysis to this area (other areas like cytoplasm were not taken into account by the algorithm). Fibrillarin areas were segmented using the building block *Find Region*, selecting the signal above a visually determined threshold with the option *Custom Threshold*. The area of these zones was calculated with *Calculate Morphology Propertie*s and zones with smaller area than 20 pixels were removed from subsequent analysis with *Select Population*. Colocalization of PIP2 spots in Fibrillarin Regions was determined with *Find Position Properties*, selecting the *Cross Population* option. The PIP2 Fibrillarin-like areas were determined by first identifying the PIP2 spots as previously described. Second, the small and intense spots were selected and their area was extended by 2 pixels. Finally, the touching spots were merged as a single zone and zones bigger than 20 pixels remained segmented. Furthermore, a ring region of 2 pixels thickness was determined with Find Region, selecting as population the PIP2 Fibrillarin-like areas.

### Live cell imaging

Cells were seeded in transparent, flat-bottom 96-or 384-well plates (Revvity; #6055302, #6057302) and allowed to incubate for at least 18 hours before starting the acquisition. For experiments involving DMSO or drug treatments, the compounds were added immediately prior to the start of data acquisition. Images were captured using the Operetta High Content Screening System (PerkinElmer) with the 20X objective in either non-confocal or confocal mode. Images were quantified using the Harmony software (version 4.9) and the results analyzed with R programming environment (v 4.4).

### Drug treatments

Cells were cultured for 24 hours before being treated with DMSO (PanReac AppliChem; #A3672) or specific compounds, including Nocodazole (MedChem Express; #HY-13520), RO-3306 (MedChem Express; #HY-12529), ISA-2011B (MedChem Express; #HY-16937), and U-73122 (MedChem Express; #HY-13419), for the durations and concentrations specified in the manuscript. DMSO controls were prepared using the same volume and dilution as the corresponding drug treatments in each experiment.

### Chromatome

One million cells were initially lysed in 0.5% CHAPS (3-cholamidopropyl dimethylammonio 1-propanesulfonate) (Roche; #10810118001) in PBS (1x) for a period of 20 minutes, with the objective of disrupting the cytosolic membrane. Following this, the lysate was subjected to centrifugation for a duration of 5 minutes at 720g at a temperature of 4°C. The supernatant was harvested as a cytosolic fraction, while the nuclear pellet was resuspended in Cytoplasmic Lysis Buffer (comprising IGEPAL 0.1%, NaCl 150 mM, Tris-HCl 10 mM, pH 7, in H₂O). The sorted cells from each phase, with the exception of the mitotic phase, were placed on top of a sucrose gradient buffer (NaCl 150 mM, sucrose 25%, Tris-HCl 10 mM pH 7 in H2O) and subjected to centrifugation for a period of five minutes at a speed of 1200g at a temperature of 4°C. The purified nuclei or lysed mitotic cells were then washed three times by resuspension in Nuclei Washing Buffer (EDTA 1 mM, IGEPAL 0.1% in PBS) and centrifugation for 5 min at 1200g at 4°C. Subsequently, the pellet was resuspended in Nuclei Resuspension Buffer (EDTA 1 mM, NaCl 75 mM, 50% sucrose, Tris-HCl 20 mM pH 8 in H2O), and the nuclear membrane was lysed by adding Nuclei Lysis Buffer (EDTA 0.2 mM, HEPES 20 mM pH 7.5, IGEPAL 0.1%, NaCl 300 mM in H2O), vortexing, and incubating for 5 minutes. Following a 2-minute centrifugation at 16,000 rpm at 4°C, the supernatant was harvested as the nucleoplasmic fraction, while the resulting chromatin pellet was resuspended in Benzonase Digestion Buffer (15 mM HEPES pH 7. The chromatin pellet was resuspended in 0.1% IGEPAL, 5 mg/mL TPCK, and sonicated on a Bioruptor Pico (Diagenode) for 15 cycles of 30 seconds on and 30 seconds off in 1.5 mL Diagenode tubes (Diagenode; #C30010016). Subsequently, the sonicated chromatin was digested with Benzonase enzyme (VWR; #706643; 2.5U) for 30 minutes at room temperature. The resulting sample was harvested as a Chromatome fraction. Unless otherwise stated, all steps were conducted on ice, and all buffers were supplemented with proteinase inhibitors (Roche; #4693132001). The concentrations of the cytosolic and chromatome extracts were determined using the Pierce BCA Protein Assay Kit (Thermo Scientific; #PIER23225).

### Western Blot

The samples were combined with 4X Laemmli sample buffer (Bio-Rad; #1610747) and heated to 95°C for three minutes. The proteins were separated by sodium dodecyl sulphate–polyacrylamide gel electrophoresis and transferred to a nitrocellulose membrane by wet transfer (25mM Tris-Base, 192mM Glycine, 20% methanol in H₂O). The membranes were then blocked for a minimum of 15 minutes at room temperature in 5% skim milk in 0.05% Tween 20 in PBS (1x). The membranes were incubated with the primary antibodies diluted in 0.05% Tween 20 in PBS (1x), for either 16 hours at 4°C or for one hour at room temperature. The membranes were washed 3 times with 0.05% Tween 20 in PBS (1x), the secondary antibodies, diluted in 0.05% Tween 20 in PBS (1x), were then applied for a period of 1 hour at room temperature. Finally, the membrane was washed 3 times with 0.05% Tween 20 in PBS (1x). The detection of the protein was done using the Odyssey CLx (Li-Cor) system, and the resulting data were analyzed using Image Studio Lite (version 5.2.5).

The following primary antibodies were used: Vinculin (Cell Signaling; #13901; 1:1000), H3 (Cell Signaling; #14269; 1:1000), H3S10ph (Merck; #06-507; 1:2000). The following secondary antibodies were used: Alexa Fluor 800 donkey anti-rabbit (Thermo Fisher Scientific; #A32735; 1:10000) and Alexa Fluor 680 goat anti-mouse (Thermo Fisher Scientific; #A21244; 1:10000).

### Mass Spectrometry

#### Liquid chromatography coupled to tandem mass spectrometry (LC-MS/MS)

The protein concentrations from chromatin enriched samples were determined using the BCA protein assay kit (Applichem CmBH, Darmstadt,Germany), and 10 μg per sample was processed using an adapted Single-Pot solid-phase-enhanced sample preparation (SP3) methodology^108^. Briefly, equal volumes (125 μl containing 6250 µg) of two different kind of paramagnetic carboxylate modified particles (SpeedBeads 45152105050250 and 65152105050250; GE Healthcare) were mixed, washed three times with 250 µl water and reconstituted to a final concentration of 50 μg/μl with LC-MS grade water (LiChrosolv; MERCK KgaA). Samples were filled up to 100 µL with stock solutions to reach a final concentration of 2% SDS, 100mM HEPES, pH 8.0, and proteins were reduced by incubation with a final concentration of 10 mM DTT for 1 hour at 56°C. After cooling down to room temperature, reduced cysteines were alkylated with iodoacetamide at a final concentration of 55 mM for 30 min in the dark. For tryptic digestion, 400 μg of mixed beads were added to reduced and alkylated samples, vortexed gently and incubated for 5 minutes at room temperature. The formed particles-protein complexes were precipitated by addition of acetonitrile to a final concentration of 70% [V/V], mixed briefly via pipetting before incubating for 18 minutes at room temperature. Particles were then immobilized using a magnetic rack (DynaMag-2 Magnet; Thermo Fisher Scientific) and supernatant was discarded. SDS was removed by washing two times with 200 μl 70% ethanol and one time with 180 μl 100% acetonitrile. After removal of organic solvent, particles were resuspended in 100 μl of 50 mM NH4HCO3 and samples digested by incubating with 1 μg of Trypsin overnight at 37°C. Samples were acidified to a final concentration of 1% Trifluoroacetic acid (Uvasol; MERCK KgaA) prior to immobilizing the beads on the magnetic rack. Peptides were desalted using C18 solid phase extraction spin columns (Pierce Biotechnology, Rockford, IL). Finally, eluates were dried in a vacuum concentrator and reconstituted in 10 µl of 0.1% TFA.

Mass spectrometry was performed on an Orbitrap Fusion Lumos mass spectrometer (ThermoFisher Scientific, San Jose, CA) coupled to an Dionex Ultimate 3000RSLC nano system (ThermoFisher Scientific, San Jose, CA) via nanoflex source interface. Tryptic peptides were loaded onto a trap column (Pepmap 100 5μm, 5 × 0.3 mm, ThermoFisher Scientific, San Jose, CA) at a flow rate of 10 μL/min using 0.1% TFA as loading buffer. After loading, the trap column was switched in-line with a 50 cm, 75 µm inner diameter analytical column (packed in-house with ReproSil-Pur 120 C18-AQ, 3 μm, Dr. Maisch, Ammerbuch-Entringen, Germany). Mobile-phase A consisted of 0.4% formic acid in water and mobile-phase B of 0.4% formic acid in a mix of 90% acetonitrile and 10% water. The flow rate was set to 230 nL/min and a 90 min gradient used (4 to 24% solvent B within 82 min, 24 to 36% solvent B within 8 min and, 36 to 100% solvent B within 1 min, 100% solvent B for 6 min before bringing back solvent B at 4% within 1 min and equilibrating for 18 min).

Analysis was performed in a data-independent acquisition (DIA) mode. Full MS scans were acquired with a mass range of 375 -1250 m/z in the orbitrap, an RF lens set at 40%, and at a resolution of 120,000 (at 200 m/z). The automatic gain control (AGC) was set to a target of 4 × 105, and a maximum injection time of 54 ms was applied, scanning data in profile mode. MS1 scans were followed by 41 MS2 customed windows. The MS2 scans were acquired in the Orbitrap at a resolution of 30,000 (at 200 m/z), with an AGC set to target 2 × 105, for a maximum injection time of 54 ms. Fragmentation was achieved with higher energy collision induced dissociation (HCD) at a fixed normalized collision energy (NCE) of 35%. A single lock mass at m/z 445.120024 was employed^109^. Xcalibur version 4.3.73.11 and Tune 3.4.3072.18 were used to operate the instrument.

The mass spectrometry data has been deposited to the ProteomeXchange Consortium via the PRIDE partner repository^110^ with the dataset identifier PXD055357.

#### Data Analysis

Chromatin data were analyzed using DIA-NN software^111^ which were subsequently normalized using the *normalize_vsn* and *median_normalisation* functions from the *DEP*^112^ and *proDA*^113^ packages, respectively^114^. The rest of the pipeline was followed according to the *DEP* package, with the inclusion of *impute.mi* function for protein-imputation from the *imp4p* package^115^. Relative enrichment of each sample was estimated as the relative abundances of known histone proteins in each sample^116^ and all biological enrichment performed using the *clusterProfiler* tool^117^, and regression correction was performed using the *rlm* function^118^. The profiles were sorted according to their similarities by calculating a distance matrix with the tool *dist*, with the *euclidean* method, followed by hierarchical clustering with the tool *hclust* with the *ward.D2* method, and further grouped in 6 clusters with the function *cutree*; these functions belong to the *stats* package^114^. Known subcellular localisations for proteins were obtained from the *pRoloc* R package, hyperLOPITU2OS2018^119^. Analysis was facilitated by the tidyverse^120^ and data.table^120^ collection of packages. Essentiality and drug sensitivity data were conducted by comparing chromatin abundances to gene essentialities and drug sensitivities^121^. Prediction of nuclear localisation signals was performed on the input fasta file using PredictNLS^122^. Evolutionary ages of proteins were derived from Gene-Ages^123^.

### Data analysis and statistics

All analyses were carried out in the R programming environment (v 4.4). Statistical parameters, including the exact value of biological replicates (e.g. total number of experiments), deviations, P-values and type of statistical test, are reported in the respective figure legends. Statistical analysis was performed across biological replicates by averaging the respective technical replicates where appropriate. Error bars in graphs represent the mean and standard deviation (SD) of at least three biologically independent experiments. Outliers outside 3 SD from the overall population were removed with a custom function named *remove_outliers*. Statistical significance was analyzed using non-parametric Wilcoxon test with the *wilcox_test* tool from the *rstatix* package^124^. P < 0.05 was considered significant.

## Data Availability

The raw mass spectrometry proteomics data have been deposited to the PRIDE repository^110^ with the dataset identifier PXD055357. Original codes are available at: https://github.com/SdelciLab/Cell_Cycle_Chromatome.

## Author contributions

**Antoni Gañez Zapater**: Conceptualization; formal analysis; validation; investigation; visualization; methodology; writing – original draft. **Savvas Kourtis**: Data curation; formal analysis; investigation; writing – original draft. **Maria Lorena Espinar Calvo**: Investigation; writing – original draft. **Laura García-López**: Investigation. **Laura Wiegan**: Investigation. **Maria Guirola**: investigation. **Frédéric Fontaine**: Data curation; formal analysis; investigation. **André C Müller**: Data curation; formal analysis; investigation; methodology. **Sara Sdelci**: Conceptualization; supervision; funding acquisition; visualization; methodology; writing – original draft; project administration.

## Acknowledgements

The authors would like to thank the CRG Flow Cytometry Facility (Barcelona, Spain) for the cell cycle analysis and to Dr. Queralt Tolosa and Dr. Ritobrata Ghose for their contributions to the generation and visualization of the figures. We acknowledge support of the Spanish Ministry of Science and Innovation through the Centro de Excelencia Severo Ochoa (CEX2020-001049-S, MCIN/AEI /10.13039/501100011033), and the Generalitat de Catalunya through the CERCA programme. The Sdelci lab’s contributions to this study has received funding from the European Research Council (ERC) under the European Union’s Horizon 2020 research and innovation programme (grant agreement No 852343), and from the Spanish Plan Estatal grants (Ministerio de Ciencia e Innovación, PID2019-110598GA-I00/AEI/10.13039/501100011033 and PID2022-141740NB-I00 funded by MICIU /AEI /10.13039/501100011033 / FEDER, U

