## Supplementary Material for "Nuclear metabolism oscillation during the cell cycle reveals a link between the phosphatidylinositol pathway and histone methylation"

### Supplementary Figures

**Supplementary Figure 1. Validation of the FUCCI-3 U2OS reporter cell line.** (A) Number of U2OS FUCCI-3 and FUCCI-4 (FUCCI-3 + H1-Maroon - H1M -) cells in G1, S and G2/M determined by integration of the FUCCI fluorescent markers measured by FACS. At least 3 biological replicates were analyzed per cell line. In each of the conditions, at least 47489 cells were considered and statistical analysis was performed using a t-test. (B) The illustration depicts the pipeline for the analysis of high-throughput microscopy images in the context of the cell cycle. The images of the FUCCI-3 fluorescent markers were acquired and subsequently merged, thereby facilitating the identification of all nuclei. A ring of eight pixels was drawn around the region of interest to measure the perinuclear intensity of the FUCCI fluorescent markers and the resulting value was used as a background reference. The background was subtracted from the mean intensity of the nuclei, thereby enabling the determination of the corrected level of each of the FUCCI fluorescent markers within the nucleus. The specific combinations of the corrected expression levels of these fluorescent markers facilitated the identification of the cell cycle phase. For the purposes of analysis, a gating procedure was performed using an XYZ projection (x-axis = SLBP-Turquoise2, y-axis = Clover-Geminin, z-axis plotted as a color gradient = Cdt1-mKO2), resulting in the FUCCI-3 plot representations, which will henceforth be referred to as such. (C) The three FUCCI-3 plots here represent the cell cycle integrity of the cell population fixed and stained for either Ki27, p21 or H3. (D) Immunofluorescence-based quantification of nuclear areas across cell cycle phases determined with the FUCCI-3 system of the cell population fixed and stained for H3. 3 biological replicates were analyzed (*n*G0 = 488, *n*G1 = 914, *n*S = 468, *n*G2/M = 436, *n*M = 58; outliers removed, 3 SD; unpaired two-tailed Wilcoxon test). (E) Immunofluorescence-based quantification of the integrated intensity of Ki67, p21 and H3 across cell cycle phases determined with the FUCCI-3 system. 3 biological replicates were analyzed (Ki67 IF: *n*G0 = 476, *n*G1 = 894, *n*S = 576, *n*G2/M = 371, *n*M = 51; p21 IF: *n*G0 = 556, *n*G1 = 765, *n*S = 396, *n*G2/M = 427, *n*M = 55; H3 IF: *n*G0 = 490, *n*G1 = 909, *n*S = 468, *n*G2/M = 435, *n*M = 58; outliers removed, 3 SD; unpaired two-tailed Wilcoxon test). (F) Growth curve of U2OS cells treated with DMSO (negative control), RO-3306 (4.5 μM; 60 hours) or Nocodazole (0.5 μM; 60 hours). FUCCI-3 U2OS cells were monitored for 60 hours. Nuclei were identified by the expression of FUCCI-3 fluorescent markers and counted over time to determine cell growth. Tracking was performed with a minimum of 200 cells per treatment. (G) Live cell imaging of cells treated with RO-3306 (4.5 μM) or Nocodazole (0.5 μM) and monitored for 60 hours. Changes in the number of Clover-Geminin negative (G1) and positive (S & G2) cells are shown in the ridge plot, illustrating the cell cycle distribution from the beginning of the treatment and every 6 hours. (H) FUCCI-3 plots showing cell cycle changes after treatment with DMSO, RO-3306 (4.5 μM) or Nocodazole (0.5 μM) and monitored in culture for up to 60 hours, the time point chosen for visualization. (I) Illustration of the gating strategy based on the FUCCI-3 plot representation used to sort the displayed populations. (J) Live cell imaging of sorted U2OS FUCCI-3 (as in H) at time points 0h, 20h and 40h after sorting, showing the progression of the cell cycle based on the synchronous oscillation of the FUCCI-3 colors. (K) Visual illustration of the in-house chromatome protocol showing how the different subcellular fractions are obtained. (L) Western blot validation of the cytoplasmic fraction using Vinculin as a cytoplasmic marker and H3 as a chromatin marker.

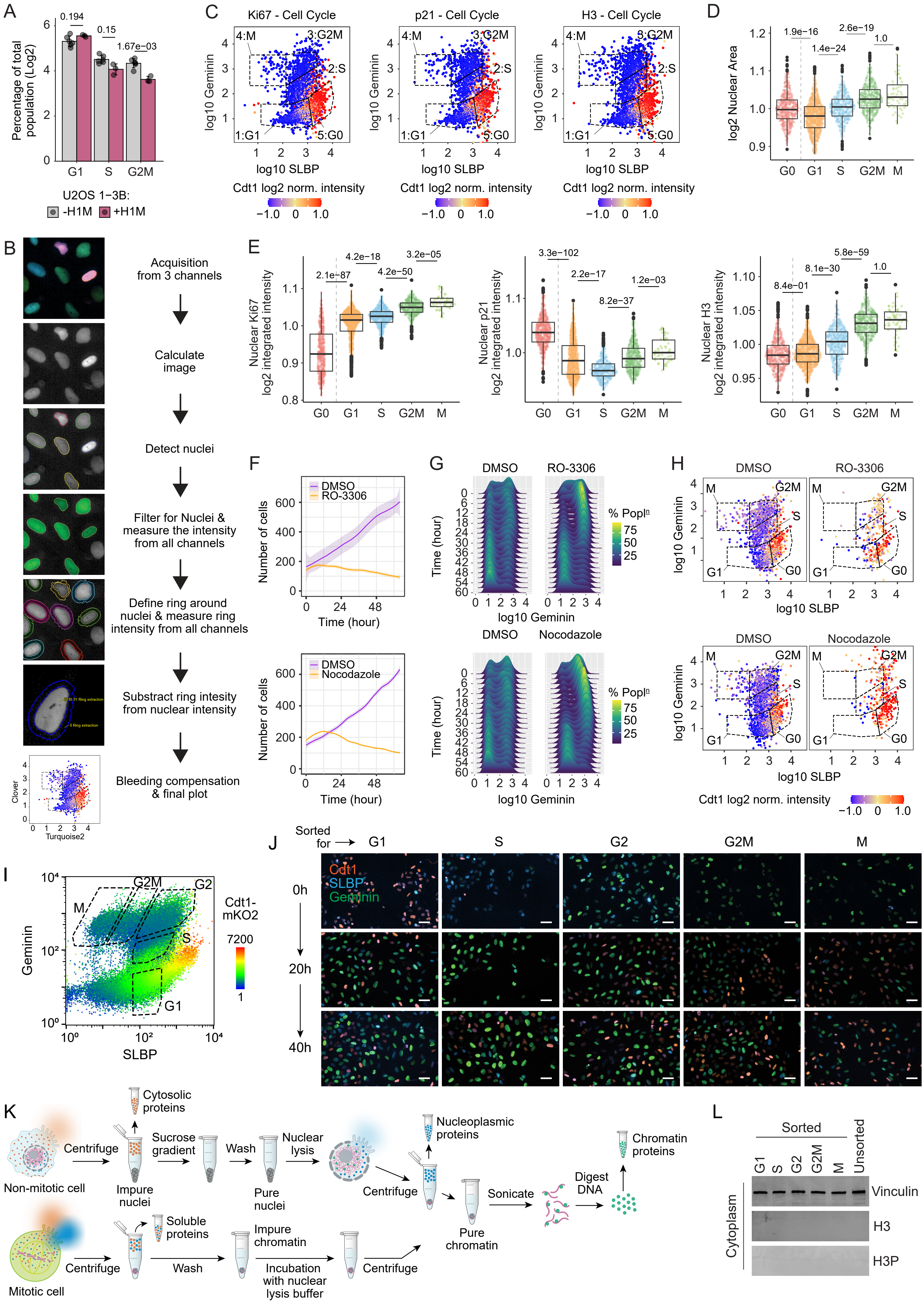
A
0.194
0.15
1.67e−03
Percentage of total population (Log2)
G1
S
G2M
U2OS 1−3B:
-H1M
+H1M
C
Ki67 - Cell Cycle
p21 - Cell Cycle
H3 - Cell Cycle
4:M
3:G2M
2:S
1:G1
5:G0
log10 Geminin
log10 SLBP
Cdt1 log2 norm. intensity
−1.0 0 1.0
D
1.9e−16
1.4e−24
2.6e−19
1.0
log2 Nuclear Area
G0
G1
S
G2M
M
B
Acquisition from 3 channels
Calculate image
Detect nuclei
Filter for Nuclei & measure the intensity from all channels
Define ring around nuclei & measure ring intensity from all channels
Substract ring intesity from nuclear intensity
Bleeding compensation & final plot
Clover
Turquoise2
E
2.1e−87
4.2e−18
4.2e−50
3.2e−05
Nuclear Ki67 log2 integrated intensity
3.3e−102
2.2e−17
8.2e−37
1.2e−03
Nuclear p21 log2 integrated intensity
8.4e−01
8.1e−30
5.8e−59
1.0
Nuclear H3 log2 integrated intensity
F
DMSO
RO-3306
Nocodazole
Number of cells
Time (hour)
G
DMSO
RO-3306
Nocodazole
Time (hour)
log10 Geminin
% Popln
75
50
25
H
DMSO
RO-3306
Nocodazole
M
G2M
S
G1
G0
log10 Geminin
log10 SLBP
Cdt1 log2 norm. intensity
−1.0 0 1.0
I
Geminin
SLBP
M
G2M
G2
S
G1
Cdt1-mKO2
7200
1
J
Sorted for
G1
S
G2
G2M
M
Cdt1
SLBP
Geminin
0h
20h
40h
K
Cytosolic proteins
Centrifuge
Sucrose gradient
Wash
Nuclear lysis
Nucleoplasmic proteins
Non-mitotic cell
Impure nuclei
Pure nuclei
Centrifuge
Sonicate
Digest DNA
Chromatin proteins
Pure chromatin
Soluble proteins
Impure chromatin
Mitotic cell
Centrifuge
Wash
Incubation with nuclear lysis buffer
Centrifuge
L
Sorted
G1
S
G2
G2M
M
Unsorted
Vinculin
H3
H3P
Cytoplasm

**Supplementary Figure 2. Curation and validation of cell cycle chromatome data.** (A) Mass spectrometry data distribution pre (raw data) and post (normalized data) normalization. Normalization of input levels of sample material was performed using the *vsn* and *median_normalisation* functions of the DEP R package[112]. (B) PCA plots (1 vs 2 and 3 vs 4) of mass spectrometry data of chromatome cell cycle samples and chromatomes from unsorted/unsynchronized cells. PCA plots were generated using the DEP R package[112] excluding any proteins which contained missing values. (C) Enrichment of known proteins to be present at different subcellular compartments in the different cell cycle phases was based on the U2OS hyperLOPIT annotations[126] (D) Clustering of proteins found to significantly change their levels of chromatin, between at least two consecutive phases, was performed using the CORREP R package[127]. (E) Clustered heatmap of proteins significantly changing chromatin abundance across phases. (F) Significantly enriched GeneOntology terms the different clusters, against a background of all chromatin detected proteins.

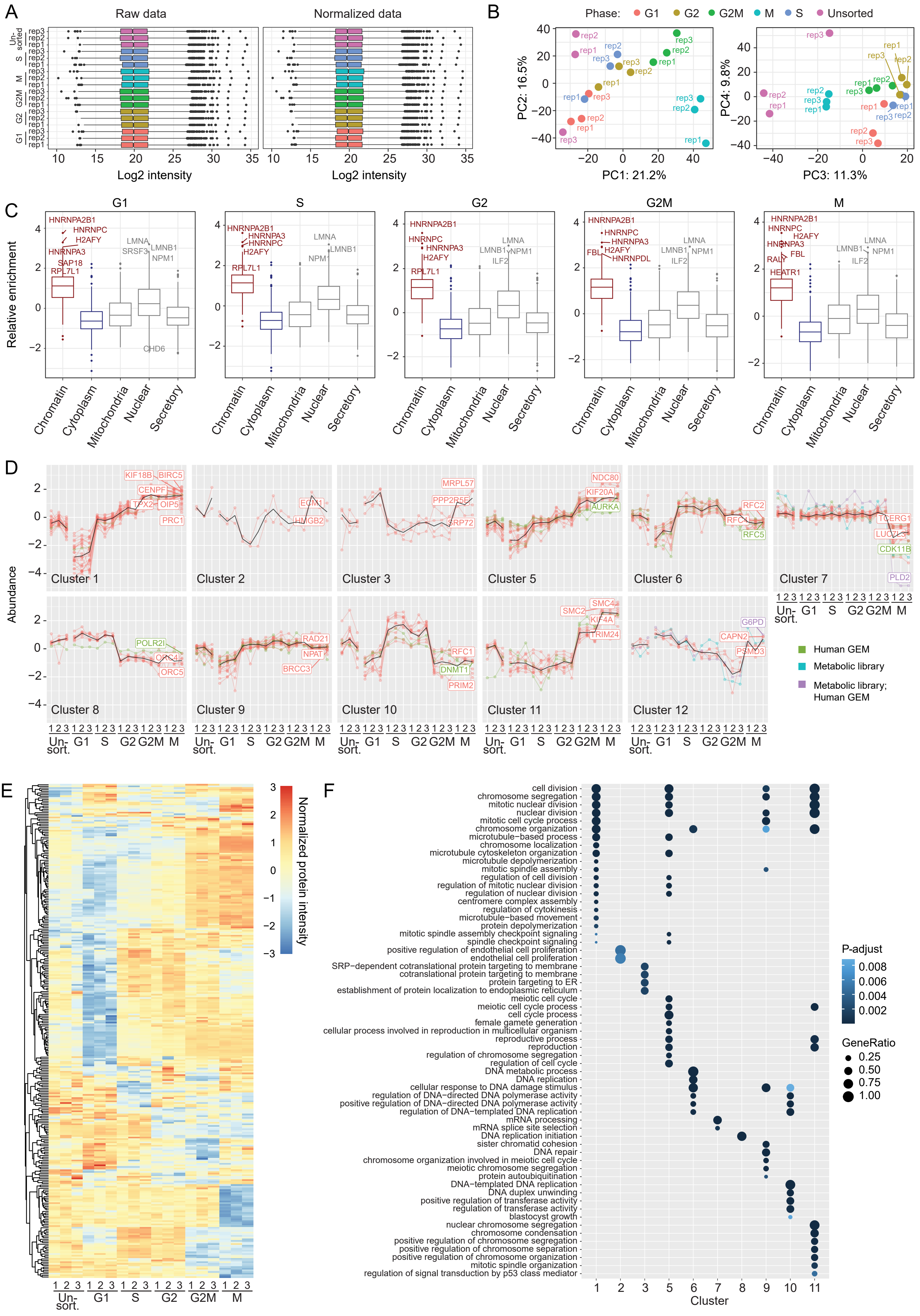

A
Raw data
Normalized data
Log2 intensity
Un-sorted
S
M
G2M
G2
G1
rep3
rep2
rep1
B
Phase: G1 G2 G2M M S Unsorted
PC1: 21.2%
PC2: 16.5%
PC3: 11.3%
PC4: 9.8%
C
G1
S
G2
G2M
M
Relative enrichment
Chromatin
Cytoplasm
Mitochondria
Nuclear
Secretory
HNRNPA2B1
HNRNPC
H2AFY
HNRNPA3
SAP18
RPL7L1
LMNA
SRSF3
LMNB1
NPM1
CHD6
ILF2
FBL
HNRNPDL
RALY
HEATR1
D
Abundance
Cluster 1
Cluster 2
Cluster 3
Cluster 5
Cluster 6
Cluster 7
Cluster 8
Cluster 9
Cluster 10
Cluster 11
Cluster 12
KIF18B
BIRC5
CENPF
TPX2
OIP5
PRC1
ECM1
HMGB2
MRPL57
PPP2R5E
SRP72
NDC80
KIF20A
AURKA
RFC2
RFC4
RFC5
TCERG1
LUC7L3
CDK11B
PLD2
POLR2I
ORC4
ORC5
RAD21
NPAT
BRCC3
RFC1
DNMT1
PRIM2
SMC2
SMC4
KIF4A
TRIM24
G6PD
CAPN2
PSMD3
Human GEM
Metabolic library
Metabolic library; Human GEM
Un-sort.
E
Normalized protein intensity
F
cell division
chromosome segregation
mitotic nuclear division
nuclear division
mitotic cell cycle process
chromosome organization
microtubule−based process
chromosome localization
microtubule cytoskeleton organization
microtubule depolymerization
mitotic spindle assembly
regulation of cell division
regulation of mitotic nuclear division
regulation of nuclear division
centromere complex assembly
regulation of cytokinesis
microtubule−based movement
protein depolymerization
mitotic spindle assembly checkpoint signaling
spindle checkpoint signaling
positive regulation of endothelial cell proliferation
endothelial cell proliferation
SRP−dependent cotranslational protein targeting to membrane
cotranslational protein targeting to membrane
protein targeting to ER
establishment of protein localization to endoplasmic reticulum
meiotic cell cycle
meiotic cell cycle process
cell cycle process
female gamete generation
cellular process involved in reproduction in multicellular organism
reproductive process
reproduction
regulation of chromosome segregation
regulation of cell cycle
DNA metabolic process
DNA replication
cellular response to DNA damage stimulus
regulation of DNA−directed DNA polymerase activity
positive regulation of DNA−directed DNA polymerase activity
regulation of DNA−templated DNA replication
mRNA processing
mRNA splice site selection
DNA replication initiation
sister chromatid cohesion
DNA repair
chromosome organization involved in meiotic cell cycle
meiotic chromosome segregation
protein autoubiquitination
DNA−templated DNA replication
DNA duplex unwinding
positive regulation of transferase activity
regulation of transferase activity
blastocyst growth
nuclear chromosome segregation
chromosome condensation
positive regulation of chromosome segregation
positive regulation of chromosome separation
positive regulation of chromosome organization
mitotic spindle organization
regulation of signal transduction by p53 class mediator
Cluster
P-adjust
GeneRatio

**Supplementary Figure 3. Metabolic enzymes oscillating on chromatin during the cell cycle.** (A) Relative chromatin abundance of metabolism-related proteins found in association with chromatin in a cell cycle dependent manner in our mass spectrometry analysis of phase-specific chromatin fractions. Magenta lines mean that the difference between the 2 consecutive phases was significant. Black lines mean that no significant difference was found between consecutive phases. Semi-transparent dots mean that the value was imputed. Statistical analysis was performed as in Figure S2B. The proteins were sorted using a hierarchical clustering algorithm based on similarities of protein abundance on chromatin across the cell cycle, and grouped in 6 different clusters, shown here in different colors. (B) FUCCI-3 plots showing the integrity of the cell cycle in the cell population used for the immunofluorescence detection of DNMT1, (C) KMT5A and (D) KDM5B. (E) Immunofluorescence-based quantification of the integrated intensity of DNMT1, (F) KMT5A and (G) KDM5B across cell cycle phases determined with the FUCCI-3 system.

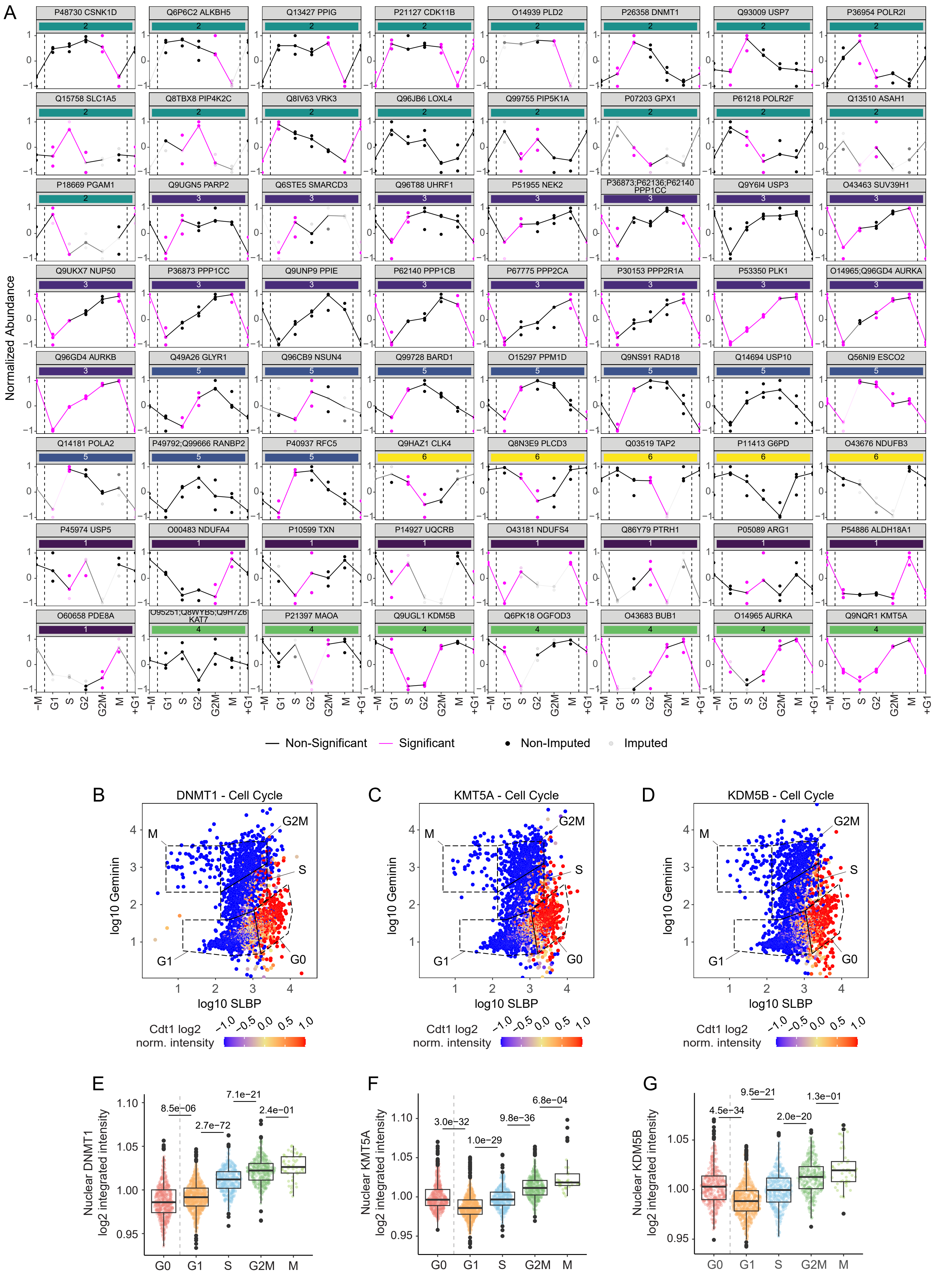
A
P48730 CSNK1D
Q6P6C2 ALKBH5
Q13427 PPIG
P21127 CDK11B
O14939 PLD2
P26358 DNMT1
Q93009 USP7
P36954 POLR2I
Q15758 SLC1A5
Q8TBX8 PIP4K2C
Q8IV63 VRK3
Q96JB6 LOXL4
Q99755 PIP5K1A
P07203 GPX1
P61218 POLR2F
Q13510 ASAH1
P18669 PGAM1
Q9UGN5 PARP2
Q6STE5 SMARCD3
Q96T88 UHRF1
P51955 NEK2
P36873;P62136;P62140 PPP1CC
Q9Y6I4 USP3
O43463 SUV39H1
Q9UKX7 NUP50
P36873 PPP1CC
Q9UNP9 PPIE
P62140 PPP1CB
P67775 PPP2CA
P30153 PPP2R1A
P53350 PLK1
O14965;Q96GD4 AURKA
Q96GD4 AURKB
Q49A26 GLYR1
Q96CB9 NSUN4
Q99728 BARD1
O15297 PPM1D
Q9NS91 RAD18
Q14694 USP10
Q56NI9 ESCO2
Q14181 POLA2
P49792;Q99666 RANBP2
P40937 RFC5
Q9HAZ1 CLK4
Q8N3E9 PLCD3
Q03519 TAP2
P11413 G6PD
O43676 NDUFB3
P45974 USP5
O00483 NDUFA4
P10599 TXN
P14927 UQCRB
O43181 NDUFS4
Q86Y79 PTRH1
P05089 ARG1
P54886 ALDH18A1
O60658 PDE8A
O95251;Q8WYB5;Q9H7Z6 KAT7
P21397 MAOA
Q9UGL1 KDM5B
Q6PK18 OGFOD3
O43683 BUB1
O14965 AURKA
Q9NQR1 KMT5A
Normalized Abundance
−M G1 S G2 G2M M +G1
Non-Significant
Significant
Non-Imputed
Imputed
B
DNMT1 - Cell Cycle
C
KMT5A - Cell Cycle
D
KDM5B - Cell Cycle
M
G2M
S
G1
G0
log10 Geminin
log10 SLBP
Cdt1 log2 norm. intensity
−1.0 −0.5 0.0 0.5 1.0
E
Nuclear DNMT1 log2 integrated intensity
8.5e−06
2.7e−72
7.1e−21
2.4e−01
F
Nuclear KMT5A log2 integrated intensity
3.0e−32
1.0e−29
9.8e−36
6.8e−04
G
Nuclear KDM5B log2 integrated intensity
4.5e−34
9.5e−21
2.0e−20
1.3e−01
G0 G1 S G2M M

**Supplementary Figure 4. Analysis of PIP2 metabolism in the nucleus.** (A) FUCCI-3 plot representing the cell cycle integrity of the cell population fixed and stained for PIP5K1A quantification. (B) Representation of PIP5K1A nuclear intensity measured by immunofluorescence and plotted accordingly to FUCCI-3-determined cell cycle phases. (C) Immunofluorescence-based quantification of the integrated intensity of PIP5K1A across cell cycle phases determined with the FUCCI-3 system. 3 biological replicates were analyzed (*n*G0 = 533, *n*G1 = 635, *n*S = 381, *n*G2/M = 331, *n*M = 52; outliers removed, 3 SD; unpaired two-tailed Wilcoxon test). (D) FUCCI-3 plot representing the cell cycle integrity of the cell population fixed and stained for PLCD3 quantification. (E) Representation of PLCD3 nuclear intensity measured by immunofluorescence and plotted accordingly to FUCCI-3-determined cell cycle phases. (F) Immunofluorescence-based quantification of the integrated intensity of PLCD3 across cell cycle phases determined with the FUCCI-3 system. 3 biological replicates were analyzed (*n*G0 = 435, *n*G1 = 690, *n*S = 445, *n*G2/M = 368, *n*M = 49; outliers removed, 3 SD; unpaired two-tailed Wilcoxon test). (G) FUCCI-3 plot representing the cell cycle integrity of the cell population fixed and stained for PLD2 quantification. (H) Representation of PLD2 nuclear intensity measured by immunofluorescence and plotted accordingly to FUCCI-3-determined cell cycle phases. (I) Immunofluorescence-based quantification of the integrated intensity of PLD2 across cell cycle phases determined with the FUCCI-3 system. 3 biological replicates were analyzed (*n*G0 = 478, *n*G1 = 754, *n*S = 422, *n*G2/M = 345, *n*M = 34; outliers removed, 3 SD; unpaired two-tailed Wilcoxon test). (J) FUCCI-3 plot representing the cell cycle integrity of the cell population fixed and stained for PIP2 quantification. (K) Representation of PIP2 nuclear intensity measured by immunofluorescence and plotted accordingly to FUCCI-3-determined cell cycle phases. (L) Immunofluorescence-based quantification of the integrated intensity of PLD2 across cell cycle phases determined with the FUCCI-3 system. 3 biological replicates were analyzed (*n*G0 = 513, *n*G1 = 727, *n*S = 408, *n*G2/M = 349, *n*M = 44; outliers removed, 3 SD; unpaired two-tailed Wilcoxon test).

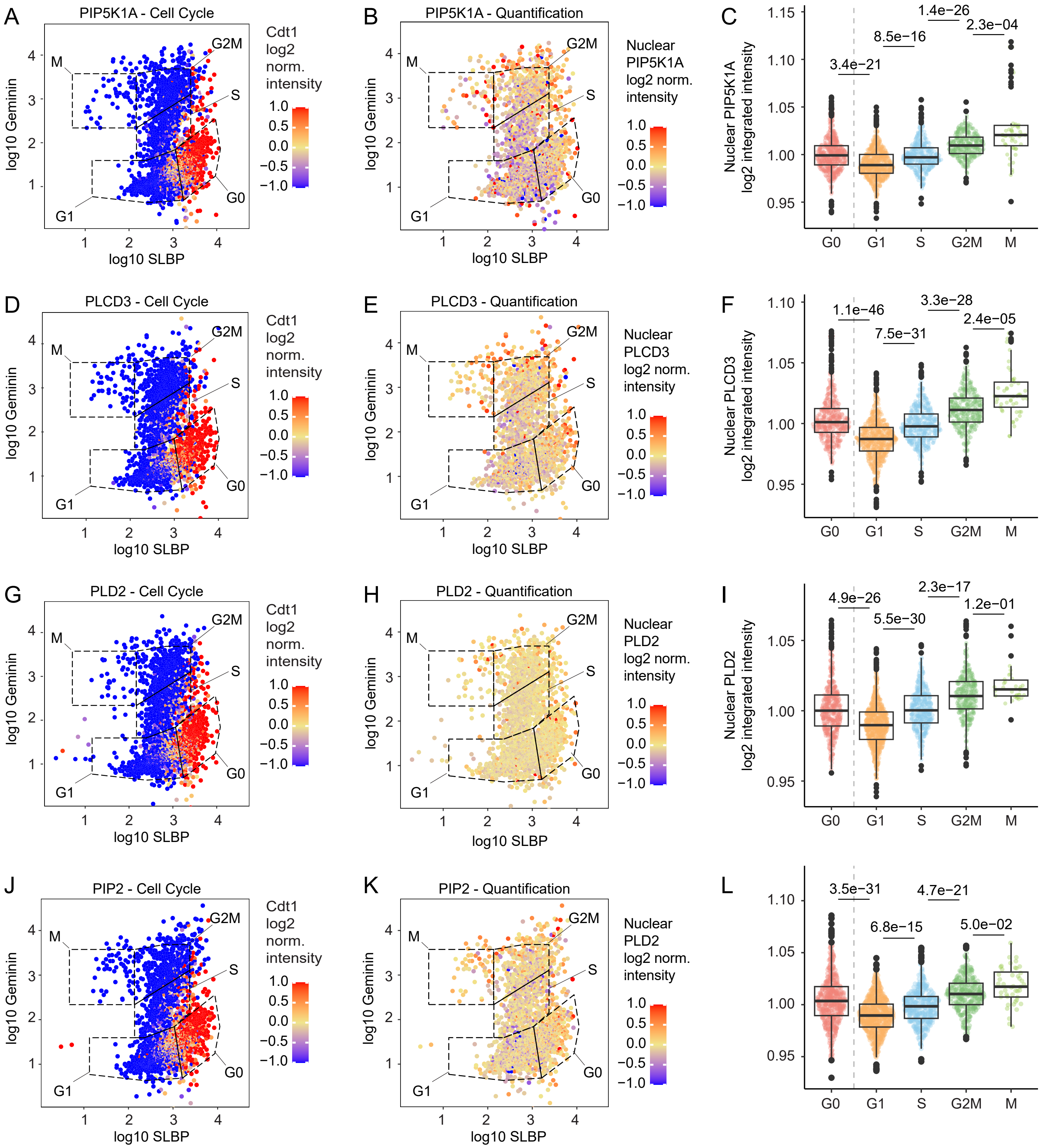
A
PIP5K1A - Cell Cycle
log10 Geminin
log10 SLBP
Cdt1 log2 norm. intensity
B
PIP5K1A - Quantification
Nuclear PIP5K1A log2 norm. intensity
C
Nuclear PIP5K1A log2 integrated intensity
3.4e−21
8.5e−16
1.4e−26
2.3e−04
D
PLCD3 - Cell Cycle
E
PLCD3 - Quantification
Nuclear PLCD3 log2 norm. intensity
F
Nuclear PLCD3 log2 integrated intensity
1.1e−46
7.5e−31
3.3e−28
2.4e−05
G
PLD2 - Cell Cycle
H
PLD2 - Quantification
Nuclear PLD2 log2 norm. intensity
I
Nuclear PLD2 log2 integrated intensity
4.9e−26
5.5e−30
2.3e−17
1.2e−01
J
PIP2 - Cell Cycle
K
PIP2 - Quantification
Nuclear PLD2 log2 norm. intensity
L
3.5e−31
6.8e−15
4.7e−21
5.0e−02
G0
G1
S
G2M
M

**Supplementary Figure 5. Nucleoli alteration following PIP2 metabolism perturbation.** (A) Example of Fibrillarin segmentation (shown in green) and the colocalization of PIP2 foci inside Fibrillarin regions (shown in red). The nucleus is shown in blue via Hoechst staining; scale bar is 10µm. (B) Immunofluorescence-based quantification of the area-related changes of nuclear Fibrillarin (nucleoli) or non-Fibrillarin (rest of the nucleus) regions in different cell cycle phases determined with the FUCCI-3 system. 3 biological replicates were analyzed (*n*G0 = 713, *n*G1 = 848, *n*S = 394, *n*G2/M = 399, *n*M = 36; outliers removed, 3 SD; unpaired two-tailed Wilcoxon test). (C) Immunofluorescence-based quantification of the intensity of Fibrillarin in Fibrillarin regions (nucleoli) in different cell cycle phases determined with the FUCCI-3 system. 3 biological replicates were analyzed (*n*G0 = 723, *n*G1 = 859, *n*S = 399, *n*G2/M = 403, *n*M = 37; outliers removed, 3 SD; unpaired two-tailed Wilcoxon test). (D) representative images showing the Fibrillarin staining in the nucleus during cell cycle progression, as well as the corresponding signal of the FUCCI-3 fluorescent marker of the depicted cell, confirming the assigned cell cycle phase. (E) Example of detection of cell cycle profiles by separating cells by Hoechst integrated intensity. (F) Dose-response curve of U2OS cells treated with U-73122 (PLCD3 inhibitor) or ISA-2011B (PIP5K1A inhibitor) for 48h to determine the IC50 of both compounds. (G) FUCCI-3 plot representing the cell cycle alterations of cells treated with U-73122 (PLCD3 inhibitor) or ISA-2011B (PIP5K1A inhibitor) for 60h or (H) 48 hours. (I) Representation of PLCD3 nuclear intensity measured by immunofluorescence and plotted accordingly to FUCCI-3-determined cell cycle phases in cells treated either with DMSO or U-73122 (5.54 µM). (J) Immunofluorescence-based quantification of the nuclear intensity of PLCD3 across cell cycle phases determined with the FUCCI-3 system in cells treated with DMSO or U-73122 (5.54 µM). 3 biological replicates were analyzed (*n*DMSO_G0 = 1182, *n*U-73122_G0 = 1085, *n*DMSO_G1 = 1845, *n*U-73122_G1 = 1048, *n*DMSO_S = 1067, *n*U-73122 = 593, *n*DMSO_G2M = 796, *n*U-73122_G2M = 522, *n*DMSO_M = 38, ISA-2011B_M = 25; outliers removed, 3 SD; unpaired two-tailed Wilcoxon test). (K) Representative images showing the changes of PLCD3 in the nucleus during cell cycle progression, as well as the corresponding signal of the FUCCI-3 fluorescent marker of the depicted cell, confirming the assigned cell cycle phase in cells treated with DMSO or U-73122 (5.54

μM). (L) Representation of PIP5K1A nuclear intensity measured by immunofluorescence and plotted according to FUCCI-3-determined cell cycle phases in cells treated either with DMSO or ISA-2011B (41.33 μM). (M) Immunofluorescence-based quantification of the nuclear intensity of PIP5K1A across cell cycle phases determined with the FUCCI-3 system in cells treated with DMSO or ISA-2011B (41.33 μM).3 biological replicates were analyzed (*n*DMSO_G0 = 906, *n*ISA-2011B_G0 = 1461, *n*DMSO_G1 = 2082, *n*ISA-2011B_G1 = 1662, *n*DMSO_S = 1356, *n*ISA-2011B = 721, *n*DMSO_G2M = 877, *n*ISA-2011B_G2M = 622, *n*DMSO_M = 107, *n*ISA-2011B_M = 33; outliers removed, 3 SD; unpaired two-tailed Wilcoxon test). (N) Representative images showing the changes of PIP5K1A in the nucleus during cell cycle progression, as well as the corresponding signal of the FUCCI-3 fluorescent marker of the depicted cell, confirming the assigned cell cycle phase in cells treated with DMSO or ISA-2011B (41.33 μM).

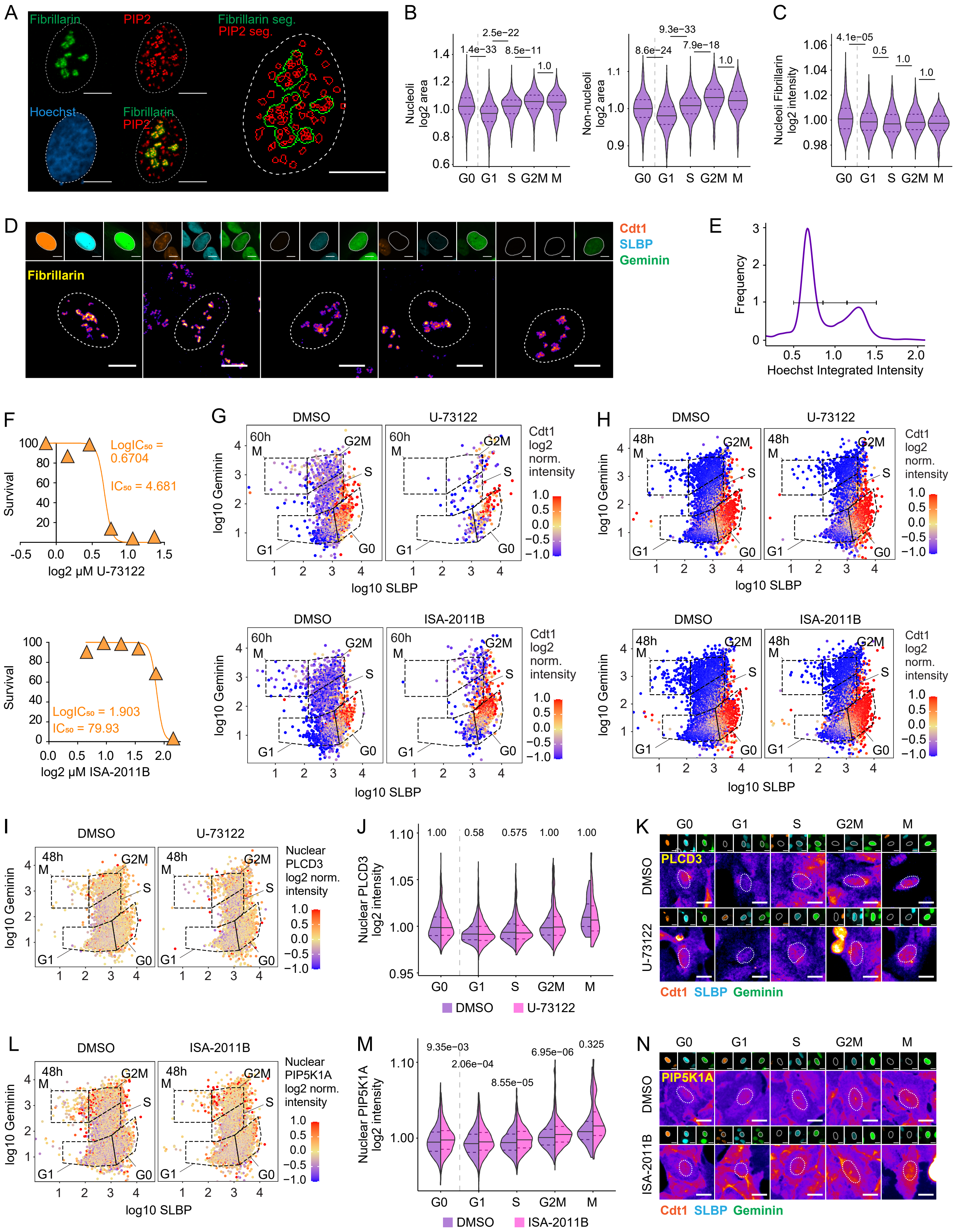

A
Fibrillarin
PIP2
Fibrillarin seg.
PIP2 seg.
Hoechst
Fibrillarin
PIP2
B
Nucleoli log2 area
Non-nucleoli log2 area
G0 G1 S G2M M
C
Nucleoli Fibrillarin log2 intensity
D
Cdt1
SLBP
Geminin
Fibrillarin
E
Frequency
Hoechst Integrated Intensity
F
Survival
LogIC50 = 0.6704
IC50 = 4.681
log2 μM U-73122
LogIC50 = 1.903
IC50 = 79.93
log2 μM ISA-2011B
G
DMSO
U-73122
ISA-2011B
60h
log10 Geminin
log10 SLBP
Cdt1 log2 norm. intensity
H
48h
I
Nuclear PLCD3 log2 norm. intensity
J
Nuclear PLCD3 log2 intensity
K
PLCD3
L
Nuclear PIP5K1A log2 norm. intensity
M
Nuclear PIP5K1A log2 intensity
N
PIP5K1A

**Supplementary Figure 6. Methylation defects linked to PIP2 metabolism perturbation.** (A) FUCCI-3 plot representing the cell cycle integrity of the cells treated with DMSO or U-73122 (5.54 μM) for 48 hours, fixed, and stained for H4K20me1. (B) Representation of H4K20me1 nuclear intensity measured by immunofluorescence and plotted according to FUCCI-3-determined cell cycle phases in cells treated either with DMSO or U-73122 (5.54 μM) for 48 hours. (C) FUCCI-3 plot representing the cell cycle integrity of the cells treated with DMSO or ISA-2011B (41.33 μM) for 48 hours, fixed, and stained for H4K20me1. (D) Representation of H4K20me1 nuclear intensity measured by immunofluorescence and plotted according to FUCCI-3-determined cell cycle phases in cells treated either with DMSO or ISA-2011B (41.33 μM) for 48 hours. (E) FUCCI-3 plot representing the cell cycle integrity of the cells treated with DMSO or U-73122 (5.54 μM) for 48 hours, fixed, and stained for H3K9me3. (F) Representation of H3K9me3 nuclear intensity measured by immunofluorescence and plotted according to FUCCI-3-determined cell cycle phases in cells treated either with DMSO or U-73122 (5.54 μM) for 48 hours. (G) FUCCI-3 plot representing the cell cycle integrity of the cells treated with DMSO or ISA-2011B (41.33 μM) for 48 hours, fixed, and stained for H3K9me3. (H) Representation of H3K9me3 nuclear intensity measured by immunofluorescence and plotted according to FUCCI-3-determined cell cycle phases in cells treated either with DMSO or ISA-2011B (41.33 μM) for 48 hours. (I) FUCCI-3 plot representing the cell cycle integrity of the cells treated with DMSO or U-73122 (5.54 μM) for 48 hours, fixed, and stained for H3K27me3. (K) Representation of H3K27me3 nuclear intensity measured by immunofluorescence and plotted according to FUCCI-3-determined cell cycle phases in cells treated either with DMSO or U-73122 (5.54 μM) for 48 hours. (J) FUCCI-3 plot representing the cell cycle integrity of the cells treated with DMSO or ISA-2011B (41.33 μM) for 48 hours, fixed, and stained for H3K27me3. (L) Representation of H3K27me3 nuclear intensity measured by immunofluorescence and plotted according to FUCCI-3-determined cell cycle phases in cells treated either with DMSO or ISA-2011B (41.33 μM) for 48 hours.

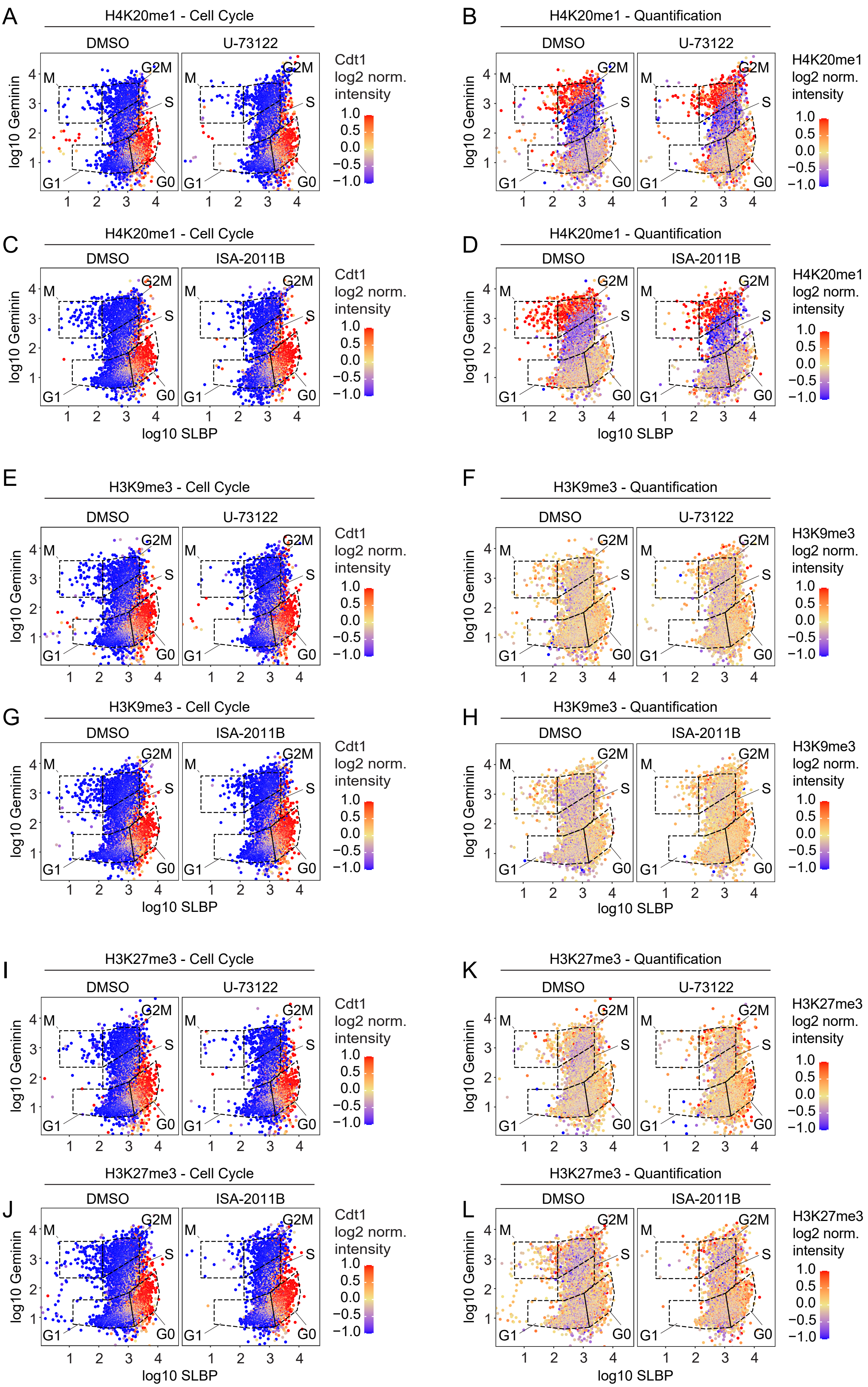
A
H4K20me1 - Cell Cycle
DMSO
U-73122
Cdt1 log2 norm. intensity
B
H4K20me1 - Quantification
H4K20me1 log2 norm. intensity
C
H4K20me1 - Cell Cycle
ISA-2011B
D
H4K20me1 - Quantification
E
H3K9me3 - Cell Cycle
F
H3K9me3 - Quantification
H3K9me3 log2 norm. intensity
G
H3K9me3 - Cell Cycle
H
H3K9me3 - Quantification
I
H3K27me3 - Cell Cycle
K
H3K27me3 - Quantification
H3K27me3 log2 norm. intensity
J
H3K27me3 - Cell Cycle
L
H3K27me3 - Quantification
M
G2M
S
G1
G0
log10 Geminin
log10 SLBP
1.0
0.5
0.0
−0.5
−1.0
